# Mosaic selection and gene flow at the onset of grain beetle domestication

**DOI:** 10.64898/2026.10.07.757314

**Authors:** Lucas Rocha Ferreira, Romuald Laso-Jadart, Leonardo Panunzi, Sana Haddad, Evangelia Eleftheriou, Benoit Vacherie, Eliaou Sellem, Karine Labadie, Thomas Lefebvre, Christophe Ambroise, Mohammed-Amin Madoui

## Abstract

Domestication is an evolutionary process that can begin before deliberate breeding, when association with humans reshapes gene flow and selection. The yellow mealworm, *Tenebrio molitor*, is undergoing this transition from a synanthropic pest of stored grain to farmed species. We asked whether gene flow between farmed populations contributes to adaptation to captivity and whether selection extends to production traits. We combined population genomics of 320 beetles from eight farmed and wild populations with genetic mapping of 15 production traits from 3,575 individuals. Farm stocks retained high diversity and weak differentiation, with Netherlands-related ancestry connecting populations across Europe and Canada. Farm-restricted sweep-like signatures were 2.06-fold enriched within regions of excess allele sharing, suggesting local selection on introgressed variation. Across 305 production-trait QTLs, overlaps with selection and introgression signals showed no significant enrichment or depletion, suggesting that adaptive gene flow may primarily contribute to adaptation to captivity rather than broadly shaping growth, development and fertility. Nevertheless, egg-hatching-rate and development-time QTLs overlapped regions of reduced nucleotide diversity and low Tajima’s *D*. These findings support a model in which gene flow supplies variation for unintended selection under intensive rearing. Crowding and physiological stress may favour adaptation to captive environments before sustained breeding establishes a broad genomic response at production loci.

## Introduction

Domestication can begin before deliberate selection for target traits, when human management changes which animals survive and reproduce^1,2^. Yet this initial phase is difficult to reconstruct because most genomic studies examine domestic animals after millennia of bottlenecks, admixtures, dispersal and breed formation^3,4^. At these endpoints, selection can appear strongly coupled to human-valued traits: long-domesticated silkworms carry extensive signatures associated with silk production and other breeding traits, and chicken sweeps overlap growth and metabolic quantitative trait loci (QTLs)^5,6^. Ancient DNA, however, showed that the derived *TSHR* allele once interpreted as central to chicken domestication was not ubiquitous in ancient European populations and probably rose during later breed formation^7^. Selection signatures in established domesticates therefore cannot automatically be attributed to the initial stages of domestication.

Animal populations sampled near the onset of human management show a more complex evolution. In the western Zagros Mountains of present-day Iran, goats managed around 10,000 years ago retained substantial genomic diversity and wild-like morphology, although sex-biased culling and restricted paternal ancestry showed human control^8^. Across the Fertile Crescent, early goat herds drew on divergent wild sources while already carrying selection signals at loci associated with pigmentation, stature, reproduction and diet^9^. These cases indicate that early domestication can alter demography, connectivity and fitness before populations lose diversity or develop domestic phenotypes.

Population genetics predicts the same separation between captive adaptation and production selection^10^. Rapid responses can draw on standing variation and involve small changes at many loci, producing weaker and less repeatable footprints than a shared hard sweep^11^. Consistent with this prediction, rabbit domestication largely shifted the frequencies of many standing variants^12^, while sustained wild–domestic gene flow in pigs coexisted with localised selection^13^. In steelhead, one captive generation produced heritable expression differences in 723 genes involved in immunity, metabolism and wound repair, despite little genome-wide differentiation^14^. Shared-origin black soldier fly lines also showed rapid population-specific differentiation and heterogeneous candidate selection signals over approximately six years^15^. High diversity and heterogeneous genomic responses to captivity therefore may characterise an early phase in which selection imposed by local captive conditions predominates over deliberate human selection for production traits.

Yellow mealworm, *Tenebrio molitor*, provides a new contemporary test of this phase. It was a long-established pest of flour and stored grain, and is thought to have originated in the eastern Mediterranean^16^. Its occurrence in Roman grain cargo and eighteenth-century stores in Québec is consistent with European dispersal through Roman grain-supply networks and later transatlantic commerce^17,18^. Commercial rearing of *T. molitor* possibly began by 1938 in the Netherlands and the 1950s in the United States, and its twenty-first-century industrial scale-up for food and feed remains recent relative to conventional livestock^19–21^. This shift towards intensive production builds on the species’ long association with human environments, imposing new selective pressures through high larval densities, standardised diets and controlled temperature and humidity^22^. In this study, we asked whether intensive rearing has produced a common genomic response at production-trait loci or whether genomic change remains structured mainly by ancestry and heterogeneous farm environments. We contrasted two non-exclusive models. A production-first model predicts repeated farm-associated signals and enrichment within QTLs for development, growth and reproduction. An ecology-and-ancestry-first model predicts retained diversity, reticulate ancestry and predominantly farm-specific selection-like signals. If movement among stocks also supplies variants, excess allele sharing should locally coincide with these signals, but neither pattern should be enriched at measured production QTLs. To test these models, we combined pooled whole-genome sequencing of 320 beetles from eight populations (40 from each of six industrial farms, one laboratory line and one synanthropic non-farmed reference) with high-throughput genotyping of 3,575 individuals at 284,452 SNPs (Extended Data Fig. 1). This design enabled QTL mapping across 15 production traits and a direct test of whether production-trait QTLs overlap genomic signatures of selection in farmed populations.

## Results

### Yellow mealworm structure and diversity in farm populations

To investigate genomic evolution during the early stages of *T. molitor* domestication, we sampled 320 individuals from six industrial farms, one laboratory line and one wild population (40 individuals per population; Fig. 1a). DNA was pooled within each population and subjected to Illumina whole-genome sequencing, and the mapping, which used a newly built genome integrating a genetic map, yielded 2,521,905 SNPs.

**Figure 1:**
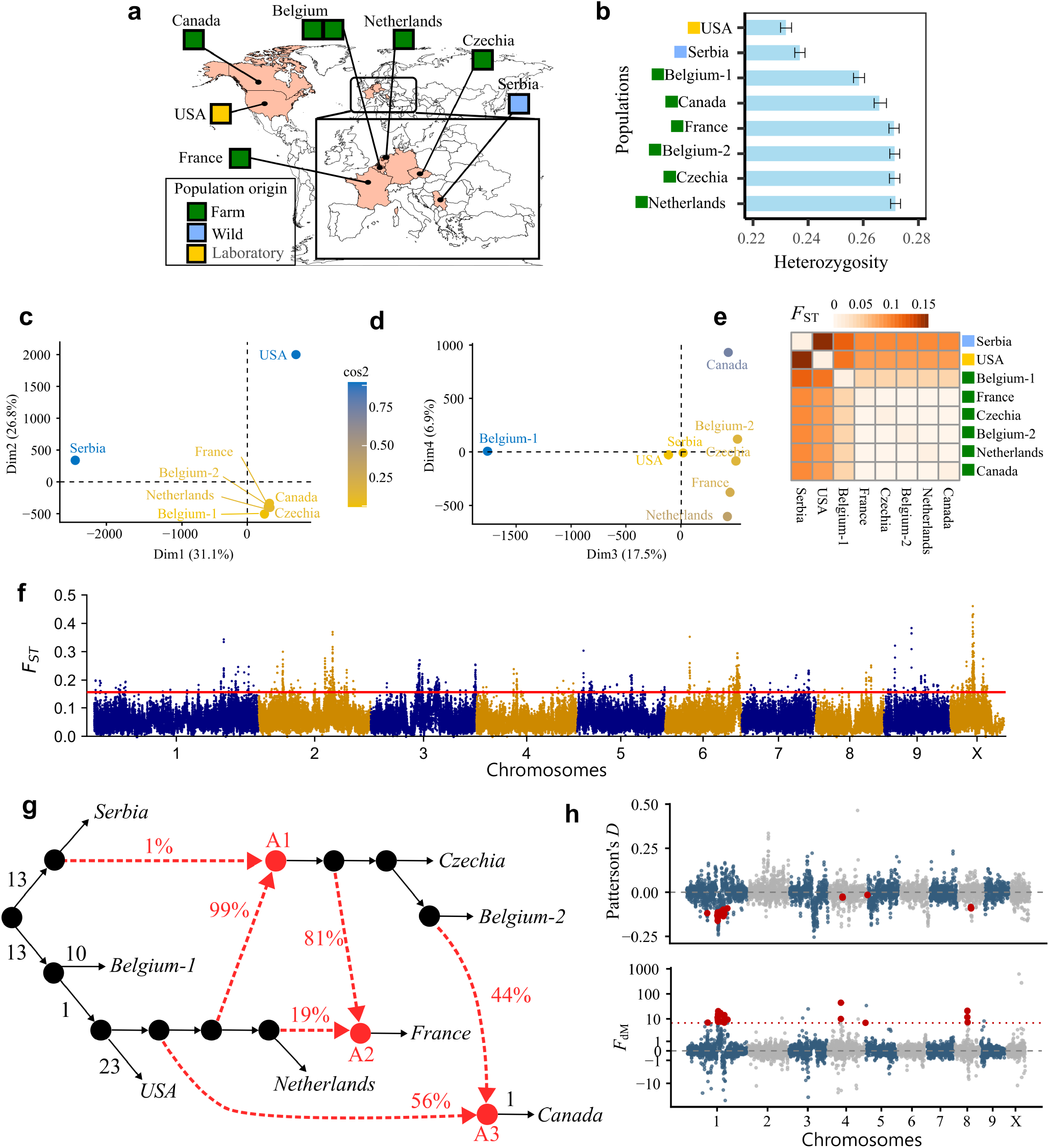
Genome diversity and population structure. **a**, Origins *Tenebrio molitor* population. **b**, Expected heterozygosity; error bars are block-jackknife standard errors. **c**,**d**, Principal-component analysis of pooled allele frequencies. **e**, Pairwise *F*_*ST*_ among populations. **f**, Genome-wide multi-population *F*_*ST*_ in overlapping 50-SNP windows; the red line marks the empirical outlier threshold. **g**, Best-fitting admixture graph inferred from *f*-statistics. Solid branches represent genetic drift and red dashed arrows indicate inferred admixture proportions. **h**, Patterson’s *D* and *F*_dM_ for *Netherlands* (P_1_), *Canada* (P_2_) and *Belgium-2* (P_3_), with *Serbia* as outgroup. Red points denote candidate introgression windows exceeding the comparison-specific *F*_dM_ threshold.

Genome-wide expected heterozygosity was higher in all six sampled farmed populations (He = 0.255–0.268) than in the *Serbia* wild population (He = 0.234; two-sided Wald Z-tests based on block-jackknife standard errors, all *P* < 0.001; Fig. 1b). Principal component analysis of allele frequencies separated the wild *Serbia* and *USA* laboratory populations from the farmed populations along the first two axes (Fig. 1c). Axes 3 and 4 further resolved fine-scale structure within the farmed group, distinguishing *Belgium-1* and *Canada* from the remaining farmed populations (Fig. 1d).

Genome-wide pairwise *F*_ST_ estimates ranged from 0.12 to 0.18 between the *Serbia* population and all other populations (Fig. 1e). Among farmed populations, mean pairwise *F*_ST_ was 0.01–0.03 and most comparisons were non-significant (permutation tests, *P* > 0.05). *Belgium-1* showed greater differentiation from the other farms (*F*_ST_ = 0.05–0.06). Although genome-wide pairwise *F*_ST_ was low among industrial populations, averaging across the genome may present localised differentiation. We therefore performed a scan of multi-population *F*_ST_ across 100,382 overlapping 50-SNP windows and identified 1,004 windows above the empirical 99th percentile (*F*_ST_ > 0.156). Merging overlapping outlier windows yielded 387 differentiated regions (Fig. 1f). Leave-one-population-out analyses identified *USA* in 170 regions, *Serbia* in 157 regions, *Belgium-1* in 59 regions, and *Netherlands* in one region as the largest contributors to regional differentiation (Extended Data Fig. 2). After excluding Wild *Serbia, USA* and *Belgium-1*, 12 regions retained at least one pairwise *F*_ST_ > 0.1 among the remaining farms, with a maximum of 0.333.

### Admixtures with *Netherlands*-related ancestry and introgressions

To place these patterns in their demographic context, we reconstructed gene flow among populations using admixture graphs. Among the tested block sizes, the best-supported graph was obtained using 20,000-SNP jackknife blocks (Extended Data Fig. 3; Extended Data Table 1). *Canada* was modelled as deriving 55.9% of its ancestry from a *Netherlands*-related ancestral branch and 44.1% from a *Belgium-2*-related branch. *France* was modelled as deriving 19.5% from the *Netherlands*-related branch and 80.5% from the lineage ancestral to *Czechia* and *Belgium-2*. This *Czechia*–*Belgium-2* ancestral lineage was itself assigned 98.6% *Netherlands*-related and 1.4% *Serbia*-related ancestry (Fig. 1g). Admixture-graph analysis therefore resolved a nested ancestry structure among the weakly differentiated farm populations, in which shared *Netherlands*-related ancestry predominated.

To determine whether the ancestry components inferred by the admixture graph were uniformly distributed across the genome, we performed a sliding-window Patterson’s *D* and *F*_dM_ analysis, retaining the *Serbia* population as the outgroup. We retained windows showing the expected negative Patterson’s *D* and falling within the comparison-specific upper 5% of positive *F*_dM_ values (*F*_dM_-5%). The genome-wide scan identified 163 candidate regions of excess allele sharing. Fifteen comparison-specific regions met the stringent criteria and were supported at multiple window scales (Fig. 1h; Extended Data Fig. 4). Six regions showed excess allele sharing between *Canada* and *Belgium-2* relative to *Netherlands*, with regional peak *F*_dM_ values ranging from 7.27 to 20.84. The strongest occurred on chromosome 8 between 10.03 and 10.18 Mb (*F*_dM_ = 20.84). In *France*, five regions showed excess sharing with the *Netherlands* relative to *Belgium-2* (peak *F*_dM_ = 3.94–9.54), and four showed excess sharing with the *Netherlands* relative to *Czechia* (peak *F*_dM_ = 6.97–26.11). The strongest signal in *France* occurred on chromosome 1 between 29.11 and 29.21 Mb indicated excess allele sharing with the *Netherlands* relative to *Czechia* (*F*_dM_ = 26.11).

### Selective-sweep signatures in farmed populations

Genome-wide Tajima’s *D* distributions differed among the eight populations (Kruskal–Wallis test, *P*<0.001). After merging adjacent windows belonging to the lower 5% of each population-specific distribution and accounting for spatial overlap with Wild *Serbia* signals, we identified 1,659 distinct low-*D*5% loci. Among them, 672 (40.5%) were restricted to farm populations, 774 (46.7%) were shared between farmed populations and *Serbia*, and 213 (12.8%) were restricted to *Serbia* (Fig. 2a). Across the six farm populations and laboratory population, 501 gene combinations occurred within loci falling simultaneously in the lowest 5% of Tajima’s *D* and nucleotide diversity in the corresponding population.

**Figure 2:**
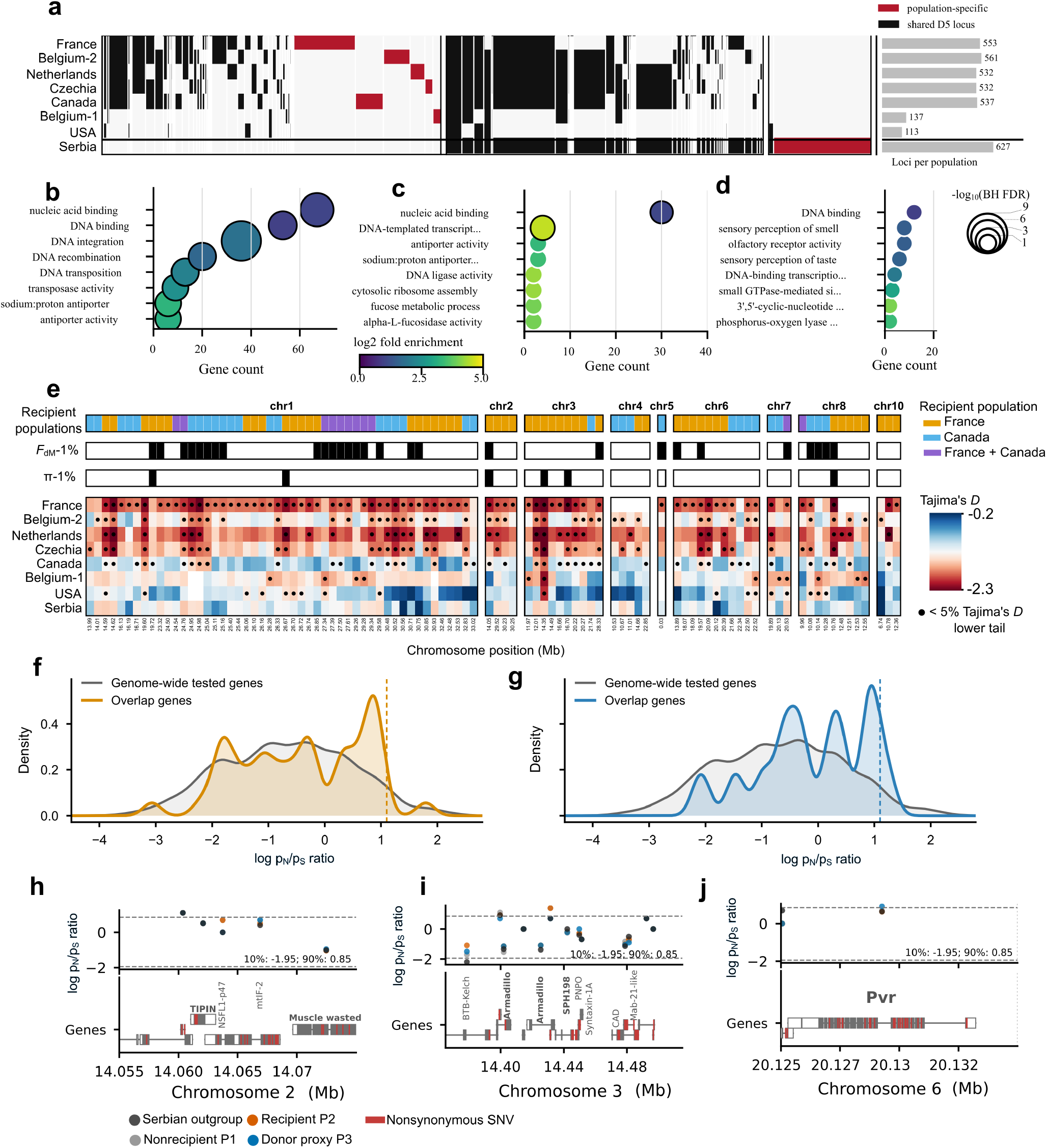
Selection signatures and adaptative introgressions. **a**, Population sharing among 1,659 merged loci in the lower 5% tail of Tajima’s *D*, classified as farm-only, shared with *Serbia* or *Serbia*-only. Bars show counts by sharing pattern and population. b–d, Gene Ontology enrichment for shared (b), farm-only (c) and Serbia-only (d) loci. Position, colour and point size represent gene count, fold enrichment and Benjamini–Hochberg FDR, respectively. e, Farm-specific low-*D* loci overlapping high-*F*_dM_ regions. Recipient populations and stricter *F*_dM_ and nucleotide-diversity thresholds are shown above the Tajima’s *D* heatmap. f,g, Distributions of log p_N_/p_S_ ratio for genome-wide and overlap genes in *France* (f) and *Canada* (g); dashed lines mark upper-5% thresholds. h–j, Population-specific coding-variation profiles for the *TIPIN/Muscle wasted* (h), *Armadillo/SPH198* (i) and *Pvr* (j) regions.

The p_N_/p_S_ log ratio distributions showed a suggestive but non-significant shift towards higher values in *Belgium-1* and *Canada* (Mann–Whitney; q=0.081), whereas no significant shift was detected in the other farms (Extended Data Fig. 5). Upper-tail enrichment was strongest in *Canada*, where 19 of 82 genes belonged to the population-specific upper 10% tail (2.31-fold enrichment; Fisher FDR=0.003). At the more stringent upper-5% threshold, enrichment was detected by the Fisher test in *Canada, Czechia* and *USA*, but these signals did not remain significant after chromosome-shift correction.

Conversely, joint low-*D*5%–low-π5% loci were not enriched significantly in the lower tail of p_N_/p_S_ log ratio in any farm after multiple-testing correction (lower-5% q≥0.913; lower-10% q≥0.113). The reduced diversity in these regions is therefore not associated with an unusual concentration of strong purifying selection on coding polymorphism. Together, the localized reductions in Tajima’s *D* and nucleotide diversity and the population-specific elevation of nonsynonymous relative to synonymous polymorphism are consistent with recent positive selection affecting a restricted subset of genes. Nevertheless, demographic events and linked background selection may also have contributed to population-specific low-diversity regions.

GO over-representation revealed distinct functional profiles among locus classes. Farm–wild shared loci were instead dominated by DNA integration, recombination, nucleic-acid binding and transposition-related annotations (Fig. 2b). Farm-only low-*D*5% loci were enriched for protein dephosphorylation, fucose metabolism, alpha-L-fucosidase and UDP-glycosyltransferase activities, suggesting potential changes in cellular signalling and carbohydrate metabolism (Fig. 2c). Wild-only loci showed weaker enrichment for cyclic-nucleotide signalling and olfactory functions (minimum q=0.099; Fig. 2d).

### Adaptive introgression in farmed populations

To identify candidate regions of adaptive introgression, we tested whether regions in the upper 5% of the positive *F*_dM_ distribution overlapped farm-specific low-D5% loci. Of 163 high-*F*_dM_5% regions, 68 overlapped farm-specific low-D5% loci (41.7%), encompassing 96 distinct low-D5% loci (Fig. 2e). This represented a 2.06-fold excess relative to randomized regions (10,000 permutations; *P*=1.0×10^−4^). We identified 5,287 allele-frequency SNV records consistent with the introgression directions within these intersections. Most were noncoding, but 198 coding SNVs corresponded to 70 unique nonsynonymous SNVs affecting 38 genes across 25 introgressed regions. Among 62 genes falling in introgression region candidates, 12 (19.4%) occurred in the upper decile of the genome-wide p_N_/p_S_ log ratio distribution, compared with 10.4% genome-wide (1.86-fold enrichment; hypergeometric *P*=0.0248; Fig. 2fg). However, none showed a significant nonsynonymous excess after multiple testing.

Genes within low-D5%–high-*F*_dM_5% overlaps were enriched for cadherin binding, calcium-dependent cysteine-type endopeptidase activity and transaminase activity, implicating cell adhesion, tissue organization, proteolysis and amino-acid metabolism. Among these loci, the Armadillo–SPH198 interval combined developmental and extracellular-proteolysis functions with direction-consistent missense variation. Two additional candidate loci contained TIPIN, a replication-fork factor potentially involved in cellular stress responses, and *Pvr*, a receptor associated with tissue homeostasis and innate immunity (Fig. 2h–j). TIPIN and *Pvr* showed direction-consistent missense variants but did not belong to the upper log-ratio tail. These loci therefore represent biologically plausible candidates for farm-associated retention of introgressed alleles, while functional and local-ancestry validation will be required to establish adaptive causality.

### Genome wide association studies on 15 production traits

The GWAS discovery population originated from a four-generation pedigreed *Tenebrio molitor* breeding population established at the Ÿnsect genetic laboratory in Évry, France, from founder stock maintained at the Ÿnsect production facility in Dole, France. The pedigree comprised 29,599 phenotyped individuals, of which 3,575 with quality-controlled genotypes constituted the genomic discovery panel. A subset of 134 individuals was sequenced at approximately 10× whole-genome coverage to discover candidate variants, which were filtered using Axiom design criteria and used to construct a custom Thermo Fisher Axiom array containing 679,205 SNPs. We screened 15 production phenotypes spanning reproduction with daily egg-laying rates (DELR), total egg count (TELC) and egg hatching rate (EHR), growth with larval mass at five ages (IMM), pupal weight (PW) and adult weight (AW) and development with egg-to-pupa (DTEP) and pupa-to-adult duration (DTPA). Association mapping of the 284,452 SNPs retained after quality control using five complementary frequentist models identified 989 significant SNP–trait associations at a 5% false-discovery rate (Fig. 3a): 216 for reproductive, 295 for growth and 478 for developmental traits. Thirteen traits yielded at least one association; no significant SNP was detected for early egg-laying rate (DELR1) and DTPA.

DTEP had the largest number of significant SNPs, followed by EHR and AW (478, 175 and 160 SNPs respectively; FDR<0.05; Fig. 3a), whereas the remaining traits each had 6–39 significant SNPs. In total, 126 SNP–trait associations significant at FDR<0.05 were supported by at least two models: 105 for DTEP, 11 for EHR, six for AW, three for late egg-laying rate and one for larval mass at day 63 (IMM63).

**Figure 3:**
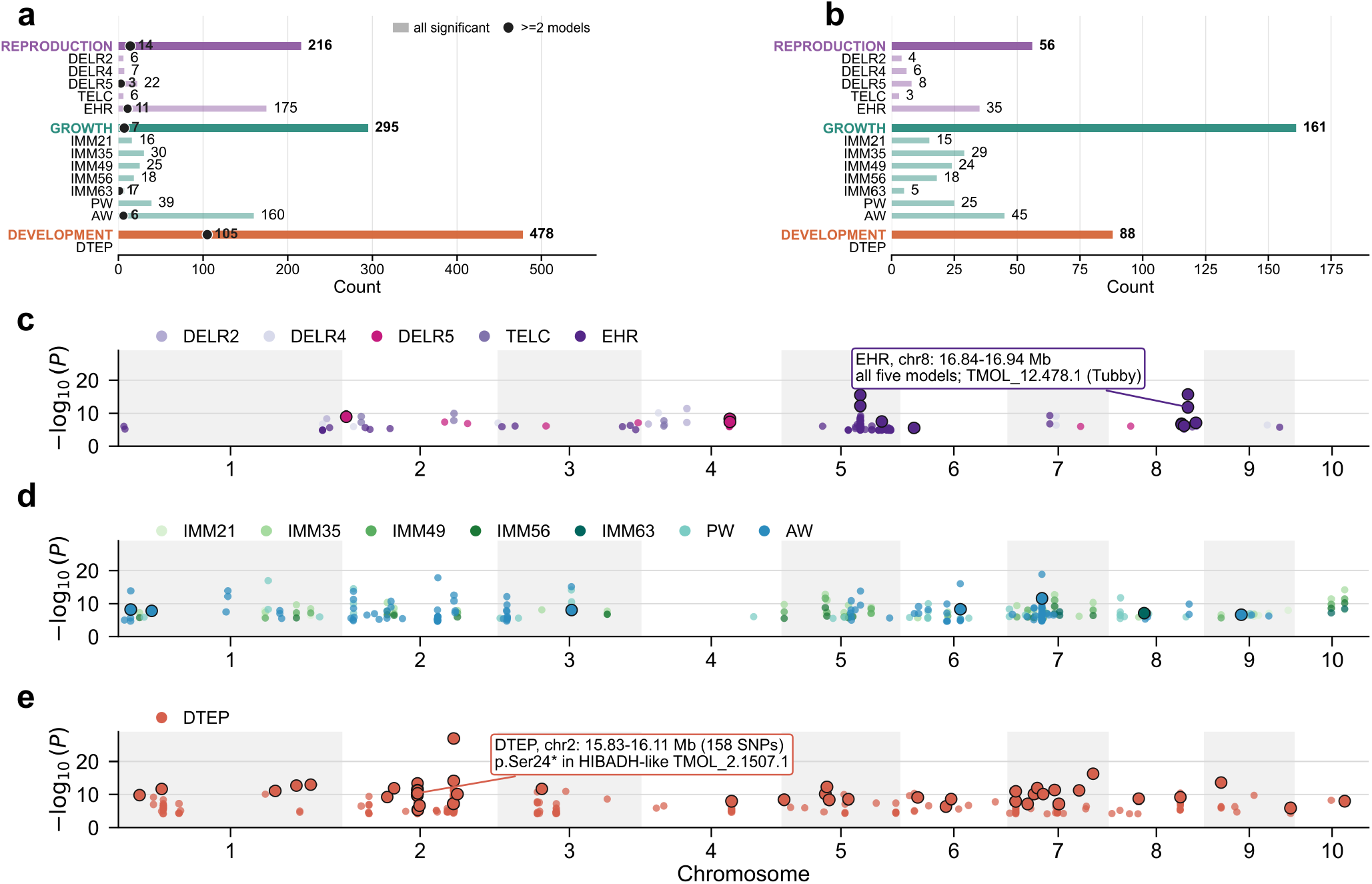
Genome-wide association landscape of production traits. **a**, Numbers of FDR-significant SNP–trait associations; black points indicate associations supported by at least two of five models. **b**, Numbers of candidate QTL intervals by trait and trait family. **c**–**e**, Positions and maximum −log_10_(*P*) of significant associations for reproductive (**c**), growth (**d**) and developmental (**e**) traits. Black outlines denote multi-model support. Callouts identify an EHR locus containing the Tubby-domain gene and a DTEP QTL containing a stop-gained variant in the HIBADH-like gene. Only significant SNPs are shown.

Based on genome-wide linkage-disequilibrium decay, we used 50 kb to define candidate QTLs, corresponding to the upper boundary of the first distance class in which mean pairwise *r*^*2*^ fell below 0.1. Significant SNPs separated by ≤50 kb were grouped, and the resulting intervals were extended by 50 kb on each side, yielding 305 candidate QTLs: 56 associated with reproductive traits, 161 with growth traits and 88 with developmental traits (Fig. 3b). As a complementary analysis, we evaluated whether DTEP-associated regions retained genomic signals in an independent validation cohort using whole-genome, robust-QTL, and reduced significant-SNP panels (Extended Data Note 1).

The genome-wide distribution of significant production-trait associations was heterogeneous, with trait-specific clusters occurring across multiple chromosomes (Fig. 3c–e). QTL analysis identified several notable candidate loci for DTEP and EHR (Extended Data Fig. 6,7 and Extended Data Note 1). An EHR locus on chromosome 8 (16.84–16.94 Mb) contained two intronic SNPs separated by <1 kb, AX-681166815 and AX-681144307, both located within a gene encoding a tubby-like protein. AX-681166815 was detected by all five association models (maximum −log_10_ *P* = 11.80), whereas AX-681144307 was supported by four models and reached a maximum −log_10_ *P* of 15.66. A DTEP interval on chromosome 2 (15.83–16.11 Mb) contained 158 SNPs significant at FDR < 0.05, including AX-674866767, a two-model stop-gained variant predicted to introduce a premature termination codon at serine residue 24 in a probable mitochondrial 3-hydroxyisobutyrate dehydrogenase-like protein.

### Production-trait QTLs show no significant enrichment in candidate selection or introgression regions

Restricting the analysis to the *French* population, that share the genetic background of the population used for QTLs detection, and accounting for uncalled genomic regions, QTL overlaps with farm-specific low Tajima’s *D* showed nominal depletion at the 10% and 5% cutoffs (*P* = 0.019 and 0.016), whereas low π showed nominal enrichment at 10% (*P* = 0.0072; Fig. 4a). Introgression overlaps were not more frequent than expected by chance, and only the 10% *D*–*F*_dM_ overlap showed nominal depletion among multipartite tests (*P* = 0.011). No overlap test remained significant after BH correction (all q = 1).

**Figure 4:**
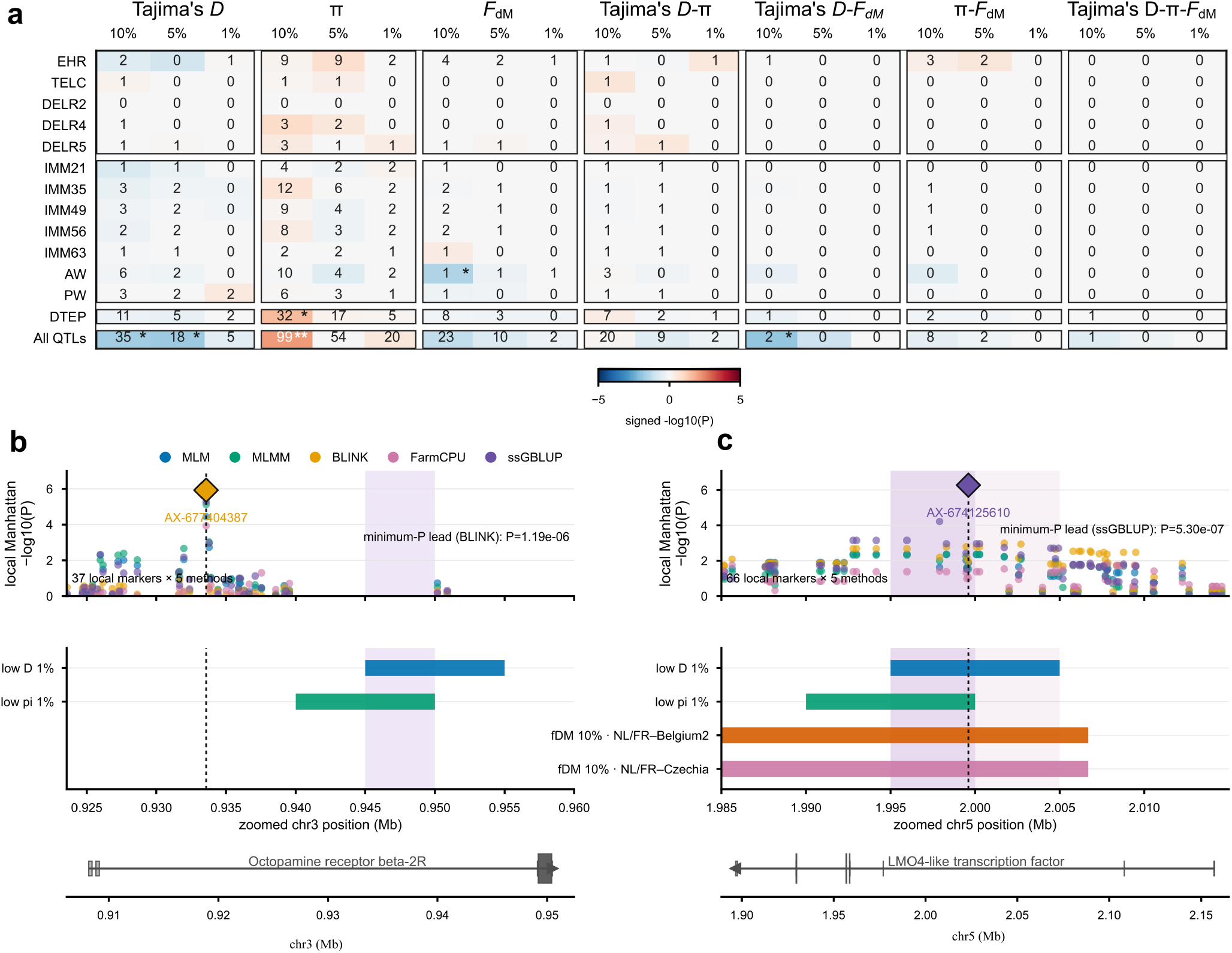
Production-trait QTL overlap with selection and introgression signals. **a**, Numbers of QTLs overlapping low Tajima’s *D*, low nucleotide diversity (π), high *F*_dM_ or their intersections at 10%, 5% and 1% thresholds. Colour gives signed −log_10_ empirical *P* from 100,000 chromosome- and QTL-width-preserving randomizations; blue and red denote depletion and enrichment. Asterisks indicate raw p-values significance (none survives BH correction). **b**, EHR QTL on chromosome 3 containing octopamine receptor beta-2R and overlapping a low-*D*/low-π segment. **c**, DTEP QTL on chromosome 5 containing the LMO4-like gene and overlapping low-*D*, low-π and high-*F*_dM_ intervals. Upper plots show association results from five models; diamonds identify lead SNPs. Lower tracks show genomic intervals and transcript structures. These examples represent locus-level convergence candidates rather than demonstrated causal colocalizations.

At the stringent 1% cutoff, overlapping farm-specific low *D* and low π identified one EHR QTL and one DTEP QTL as candidates for unintended selection on production traits. The EHR QTL (chr3:0.88–0.98 Mb) contained a ∼5-kb shared interval within an octopamine receptor β-2 gene; its lead SNP lay outside this interval but within the same transcript (Fig. 4b). The DTEP QTL (chr5:1.95–2.05 Mb) contained a ∼5-kb low-*D*–π interval encompassing its lead SNP within a gene encoding a LIM domain-only protein 4-like (LMO4-like) protein (Fig. 4c). At 10%, this same QTL contained the only exact *D*–π–*F*_dM_ overlap, spanning 10 kb and including the lead SNP.

## Discussion

Understanding the role selection in the early domestication of a species is challenging. Domestication is thought to decrease genetic diversity of the farmed populations, both through successive bottlenecks and unintended selection for adaptation to farming conditions and captivity, and human-mediated selection for production traits. Here, we showed that early mealworm domestication appears to involve redistribution of genetic variation rather than a uniform loss of diversity. Weak differentiation among farms, with their higher heterozygosity than the non-farmed reference, is consistent with interconnected breeding stocks. Gene flow can replenish variation while counteracting differentiation^23^, and international trade has already distributed closely related black soldier fly stocks across continents^24^. Transfers of mealworm larvae similarly connect distant producers and generate admixture when introduced and resident stocks interbreed. The occurrence of the ancestry component predominant in Dutch farms across geographically distant populations is consistent with a shared source in the Netherlands, an early European centre of commercial mealworm rearing^25^. Under this model, production networks shaped the distribution of genetic diversity during early domestication.

Against this connected background, heterogeneous genomic signatures of selection are consistent with adaptation to intensive rearing. Captivity can generate heritable adaptation within one generation^26,27^. In mealworms, crowding raises substrate temperature and alters development^28^, diet composition affects growth and survival^29^, and density modifies investment in immunity^30^. The enrichment for fucose metabolism and α-L-fucosidase activity among farm-only candidate loci suggests a possible contribution of dietary adaptation. α-L-fucosidases can participate in the breakdown of fucosylated plant glycans, raising the possibility that these genomic signatures reflect selection on the utilisation of complex plant carbohydrates supplied in farm diets^31,32^. Variation in husbandry could generate farm-specific selection, while polygenic adaptation to shared industrial stressors could produce different genomic responses among farms. In replicated *Drosophila* populations, similar phenotypes evolved through different combinations of alleles^33^.

Routine husbandry also provides a mechanism linking production management to reproductive success before formal breeding programmes. While no breeding program has yet been finalised in mealworm, protocols describe sieving and repeated pupal collection to establish breeding cohorts^34^. Size grading into homogeneous batches could impose selection when particular size classes contribute disproportionately to reproduction. Likewise, preferential recruitment of the earliest pupae could favour faster development and increase the contribution of early-developing families. Experimental selection on pupal size altered mealworm body mass and other life-history traits^35^, while quantitative-genetic analyses identified heritable variation and a trade-off between developmental duration and pupal weight^36^. Consequently, selecting larger individuals and recruiting earlier pupae could drive different, potentially opposing responses. Variation in these management decisions offers an explanation for heterogeneous signatures of selection among producers.

The local overlap between excess allele sharing and genomic signatures of selection suggests that stock exchange may supply variation subsequently favoured under local rearing conditions. An established example of this mechanism comes from Tibetan cattle, in which yak-derived alleles involved in the response to hypoxia contributed to adaptation to high altitude^37^. Our results suggest that exchanges among commercial mealworm stocks may play a comparable role from the earliest stages of domestication, supplying variation on which local farm conditions can act.

Production-trait QTLs showed no detectable enrichment within candidate signatures of selection or introgression. This pattern is compatible with selection on standing variation, which can leave soft sweep signatures^38^, and with polygenic responses distributed across alternative combinations of alleles^33^. It also raises the possibility that reproductive success under industrial conditions depends on competitive performance and stress tolerance not fully represented by the measured production traits. The developmental-time and egg-hatching QTLs overlapping candidate signatures of selection provide specific links between genomic change and traits potentially affected by breeding-cohort management. The octopamine receptor β2 candidate is functionally consistent with the egg-hatching-rate association, as silencing this receptor reduces egg hatchability and modulates stress resistance in the brown planthopper *Nilaparvata lugens*^39^.

Ancient DNA has transformed the reconstruction of early animal domestication^3^, but DNA preservation varies among tissues and environments, leaving the available record temporally and geographically uneven^40,41^. Moreover, production phenotypes can rarely be measured directly in sequenced archaeological specimens. By combining population genomics with QTL mapping in a recently farmed species, we may capture the relationship between population change and production-trait architecture as domestication starts. In mealworms, stock circulation and local genomic signatures of selection are already detectable, but links between selection and production traits are restricted to two candidate QTL overlaps and do not form a genome-wide pattern. Early domestication may therefore reorganise ancestry and fitness, while concomitant selection on production traits remains restricted to a small number of loci. Routine husbandry could initiate this connection by determining which individuals reproduce^34,35^, while structured breeding programmes could reinforce it by systematically controlling which variants enter the next generation.

## Conclusions

By integrating a high-density genetic map, population genomics and QTL mapping, we provide the first comprehensive view of genomic evolution in farmed *T. molitor*. Farmed stocks retain substantial diversity within a reticulate ancestry network shaped partly by adaptive introgression, while farming-associated signatures remain heterogeneous and weakly coupled to production-trait loci. This configuration reflects an early domestication phase, following synanthropic preadaptation but preceding sustained directional breeding. Yellow mealworm farming thus offers a unique opportunity to observe domesticated genomes taking form and to develop breeding strategies that combine productivity with genetic diversity, resilience and long-term sustainability.

## Materials and methods

### Insect rearing

The insects used to generate the F_1_ outcross family were reared at Ÿnsect genetics laboratory (Évry, France). One male and one female from two distinct lines maintained in panmixia for eight years in Yannick Moret’s laboratory from University of Burgundy (Dijon, France). Parents were crossed to generate the F_1_ offspring used for linkage mapping. Insects were maintained at 28 °C, 60% relative humidity and in darkness, with feed provided ad libitum and water gel supplied every three days. The 134 individuals sequenced for SNP-array development and the GWAS discovery population were sampled from the pedigreed reference population described by Sellem and colleagues^36^.

### F1 outcross sequencing and genotyping

The cross produced 101 offspring. Larvae were first ground individually using ZR BashingBead Lysis Tubes (2 mm) (ZymoResearch, CA, USA) and subsequently DNA extraction was performed using the Quick-DNA 96 Plus Kit (ZymoResearch). DNA libraries were prepared using the NEBNext® Ultra II DNA Library prep kit for Illumina (New England Biolabs, MA, USA) and sequenced on an Illumina NovaSeq6000 instrument (Illumina, San Diego, CA, USA) using 150 base-length read chemistry in paired-end mode to reach around 10x depth coverage by individual. Reads were aligned to the repeat-masked *Tenebrio molitor* reference genome with BWA-MEM v0.7.17^42^, and alignments were sorted and deduplicated with SAMtools v1.15.1^43^. Variants were called using the *mpileup* and *call* procedures in bcftools v1.15.1, after which the 103 individual VCF files were merged. SNPs were retained when at least one individual had a sequencing depth of at least three reads and the alternative allele was supported by reads from both strands. This procedure yielded 8,905,098 SNPs. Ninety-eight offspring were retained for construction of the final linkage map.

### Linkage-map construction and chromosome-scale pseudomolecules

Linkage maps were constructed with OneMap^44^. Markers were separated according to parental segregation type: 186,341 markers were heterozygous in the female and homozygous in the male, whereas 101,189 markers showed the reciprocal configuration. Markers showing significant segregation distortion were excluded. To reduce redundancy and computational burden, markers with an estimated pairwise recombination fraction of zero were collapsed into bins represented by the marker with the lowest proportion of missing data. Linkage groups were defined from two-point recombination estimates using a minimum LOD score of 3.0 and a maximum recombination fraction of 0.50. Genetic distances were calculated using the Kosambi mapping function. The final map contained approximately 283,000 SNPs distributed among ten linkage groups. These linkage groups were aligned to the scaffold-level reference assembly, and scaffolds were anchored and ordered with ALLMAPS^45^ to generate chromosome-scale pseudomolecules representing the nine autosomes and the X chromosome.

### Population sampling and pooled sequencing

The population-genomic panel comprised 320 individuals from eight *T. molitor* populations, with 40 individuals sampled per population. Six populations originated from industrial producers: Mystik Farms, Canada; Papek s.r.o., Czechia; Kingsect, Belgium; Pronutrix, Belgium; Ÿnsect, France; and Star Food, the Netherlands. These populations are hereafter referred to as *Canada, Czechia, Belgium-1, Belgium-2, France* and *Netherlands*, respectively. The *USA* laboratory population was supplied by the University of Thessaly, Greece. The Serbian population, supplied by the University of Burgundy, originated from a disused mill and was used as a synanthropic, non-farmed reference population and was referred to as *Serbia*.

Larvae were homogenized individually in 2-mm ZR BashingBead lysis tubes, and DNA was extracted using the Quick-DNA 96 Plus Kit. DNA concentrations were normalized, and equal quantities from the 40 individuals in each population were combined. Libraries were prepared with the NEBNext Ultra II DNA Library Prep Kit and sequenced as 150-bp paired-end reads on an Illumina NovaSeq 6000. Because sequencing yield differed among pools, each population was randomly downsampled with seqtk to 700 million paired reads, corresponding to 700 million reads in each mate file and matching the smallest library.

### Pool-seq processing and variant filtering

Adapters and low-quality bases were removed with fastp^46^. Reads shorter than 75 bp or containing five or more ambiguous bases were discarded. Filtered reads were aligned to the chromosome-scale *T. molitor* reference genome with BWA-MEM, and duplicate reads were removed using SAMtools markdup. Low-quality alignments were excluded before variant calling with bcftools. Reference- and alternative-allele frequencies were calculated from read counts. Variants were retained when the alternative-allele frequency was at least 0.01 in one or more populations, the read depth in each retained population was at least eight and did not exceed three standard deviations above that population’s genome-wide mean, no significant strand bias was detected, and the variant was a biallelic nuclear SNP. Mitochondrial variants were excluded. Read counts from the eight populations were imported into poolfstat v3.1.0^47^. The final population-genomic dataset was restricted to SNPs with a global minor-allele frequency of at least 0.01 and population-specific coverage between the 10th and 90th percentiles. These filters yielded 2,521,905 biallelic SNPs.

### Genetic diversity, population structure and differentiation

Within-population expected heterozygosity was estimated as 1-Q1 using compute.fstats, where Q1 is the probability that two gene copies sampled from the same population are identical in state. Standard errors were obtained by block jackknifing over consecutive blocks of 10,000 SNPs. Heterozygosity contrasts between each population and the Serbian reference were evaluated using two-sided Wald tests based on the jackknife standard errors. Population structure was assessed by centred and scaled principal component analysis of reference-allele read frequencies using FactoMineR^48^.

Genome-wide pairwise *F*_ST_ was estimated from unbiased identity-by-state probabilities using compute.fstats. Statistical support for pairwise contrasts was evaluated by permutation. Local differentiation was assessed using multi-population *F*_ST_ calculated in windows of 50 consecutive SNPs advanced by 25 SNPs. Block-jackknife estimates used blocks of 500 SNPs. Outliers were defined by the 99th percentile of the empirical window distribution, corresponding to (*F*_ST_ > 0.1568). Overlapping outlier windows were merged into differentiated regions. The contribution of each population to a differentiated region was evaluated by recalculating *F*_ST_ after sequentially excluding each population and computing Δ*F*_ST,i_ = *F*_ST_,all − *F*_ST,i_. Residual differentiation among industrial populations was examined after excluding *Serbia, USA* and *Belgium-1*.

### Admixture-graph inference and local excess allele-sharing

Allele-frequency covariance statistics *f*_2_, *f*_3_, *f*_4_, and Patterson’s *D* and their covariance matrix were estimated from Pool-seq read counts using compute.fstats in poolfstat v3.1.0. Uncertainty was assessed by block jackknifing. A base graph rooted with the Serbian reference included the differentiated *Belgium-1* and *USA* populations. *Netherlands, Czechia, France, Belgium-2* and *Canada* were then added sequentially with graph.builder, following the recorded chronology of the industrial populations. Searches retained up to 200 candidate graphs per iteration and discarded graphs more than six BIC units above the current optimum (max.heap.size = 200; heap.dbic = 6). Models within *ΔBIC<2* were considered similarly supported. Robustness was assessed using blocks of 1,000, 5,000, 10,000, 20,000 and 40,000 SNPs. The 20,000-SNP analysis was retained as the principal graph because it minimized residual misfit while retaining 116 jackknife blocks.

Graph-guided local allele-sharing tests used Serbia as the outgroup in three four-population configurations: (Netherlands – Canada – Belgium-2 – Serbia, Belgium-2– France – Netherlands – Serbia, and Czechia–France–Netherlands–Serbia, corresponding to the *P*_*1*_, *P*_*2*_, *P*_*3*_ and outgroup positions, respectively. In these orientations, negative Patterson’s *D* indicated excess allele sharing between *P*_*2*_ and *P*_*3*_. Patterson’s *D* and *F*_dM_ were calculated in 100-, 250- and 500-kb windows with 50% overlap. Windows containing fewer than 100 SNPs were excluded. Within each comparison and window size, thresholds were estimated from positive *F*_dM_ values among windows with (Patterson’s *D*<0). The upper 5% threshold was used for the broad scan and the upper 1% threshold for the stringent scan. Overlapping 100-kb candidate windows were merged within comparison and chromosome. Overlap with a candidate from the 250- or 500-kb scan was recorded as multiscale support but did not enlarge the 100-kb candidate interval.

### Diversity scans and population-specific low Tajima’s *D* loci

Tajima’s *D* and nucleotide diversity π were calculated with PoPoolation^49^ using 100-kb sliding windows and a 10-kb step (pool size = 40, minimum allele count = 2 and minimum coverage = 4). The resulting smoothed estimates were represented as 10-kb intervals for genomic-overlap analyses. Population-specific empirical lower-tail thresholds were calculated from all callable windows. The lower 5% tail was used for the principal selection analysis, whereas the lower 10% and 1% tails were evaluated in sensitivity and QTL-overlap analyses. Two distinct locus-construction procedures were used. For the descriptive farm-versus-*Serbia* classes and their GO analysis^50^, low Tajima’s *D* windows were initially classified as farm only, farm plus *Serbia*, or *Serbia* only. USA was excluded when defining these three classes because it represents a laboratory rather than a farm population. Adjacent windows were merged within class and chromosome. A nominal farm-only locus was reclassified as farm plus Serbia when any component farm window overlapped at least 50% of a *Serbia* low Tajima’s *D* window. This spatial correction generated the farm-only, farm-plus-Serbia and Serbia-only locus counts reported in the Results.

### Coding-polymorphism analyses

The Pool-seq VCF was annotated directly with SnpEff v5.4^51^ and analyses were restricted to biallelic SNVs. For population-specific coding counts, a site was considered callable when total depth was at least ten reads and both the reference and alternative alleles were supported by at least three reads. p_N_ and p_S_ were defined as the numbers of segregating nonsynonymous and synonymous coding sites, respectively, for each transcript and population. Genes containing at least five callable coding variants were considered usable. The log p_N_/p_S_ ratio, defined as log ((p_N_+1) / (p_S_+1)) was used as a descriptive statistic. Gene-specific synonymous and nonsynonymous mutational opportunities were calculated from the reference coding sequence. Distributions of the log p_N_/p_S_ ratio in joint-low Tajima’s *D* and π and background genes were compared using two-sided Mann–Whitney tests. Enrichment in population-specific upper and lower 5% and 10% tails was evaluated with one-sided Fisher exact tests. Benjamini–Hochberg correction^52^ was applied across the population tests within each tail definition.

### GO over-representation analyses

Genes physically intersecting each corrected low Tajima’s *D* locus class were used as foreground sets. The corresponding background comprised genes with callable data and at least one GO assignment. One-sided hypergeometric tests were restricted to terms represented by at least two foreground genes and five background genes and assigned to no more than 50% of the background. Benjamini–Hochberg correction was applied separately within each population or locus profile and separately for direct and propagated annotations.

### Candidate adaptive introgression

Broad candidate adaptive-introgression loci were defined by exact physical overlap between farm-only low Tajima’s *D* loci at the 5% threshold and merged 100-kb *F*_dM_ regions in the comparison-specific upper 5% tail. Enrichment was evaluated using 10,000 chromosome- and interval-length-preserving randomizations. Among genes intersecting the overlaps, enrichment in the genome-wide upper decile of log p_N_/p_S_ ratio was evaluated using a one-sided hypergeometric test. Opportunity-corrected binomial tests provided the inferential assessment of nonsynonymous excess. GO enrichment of exact-overlap genes followed the procedure described above, with false-discovery rates calculated separately for direct and propagated annotations.

### GWAS discovery population and phenotypes

Genome-wide association analyses were conducted in a four-generation pedigreed *T. molitor* population maintained at the Ÿnsect genetics laboratory in Évry, France. The population originated from 72 founders sampled from the Ÿnsect production facility in Dole, France^36^. Breeding pairs were selected using optimum-contribution procedures to limit increases in inbreeding while retaining genetic diversity. Adults and full-sib offspring were maintained at 28 °C and 60% relative humidity in darkness, with feed supplied ad libitum. Individual identity and parentage were recorded throughout the experiment. Fifteen phenotypes represented reproduction, growth and developmental timing. Reproductive traits comprised daily egg-laying rates during four one-week recording periods approximately 14, 35, 56 and 77 days after pair formation (DELR1, DELR2, DELR4 and DELR5), total egg count across these periods (TELC), and egg hatching rate (EHR), defined as the percentage of eggs producing larvae by day 21. Growth traits comprised mean individual larval mass at 21, 35, 49, 56 and 63 days after hatching (IMM21–IMM63), calculated as full-sib-group biomass divided by the number of live larvae, together with individually measured pupal weight (PW) and adult weight (AW). Developmental time from egg to pupa (DTEP) was converted to four ordered empirical-quartile classes, coded 1–4 from fastest to slowest development, to reduce calendar-related heaping caused by the absence of weekend observations. Developmental time from pupa to adult (DTPA) was measured in days.

### SNP-array construction and genotype quality control

Candidate variants were identified by approximately 10× whole-genome sequencing of 134 discovery-population individuals. Reads were aligned to the repeat-masked reference genome with BWA-MEM v0.7.17, alignments were processed with SAMtools v1.15.1, and variants were called with bcftools v1.18. Variants passing Thermo Fisher Scientific’s Axiom design criteria were used to construct a custom array containing 679,205 SNPs. Array genotypes were initially obtained for 3,862 samples. Samples showing evidence of plate contamination, genotype missingness greater than 2%, or duplication were excluded. Analyses were restricted to biallelic SNPs on the ten anchored pseudomolecules. SNPs were retained when their minor-allele frequency was at least 0.05, genotype missingness was no greater than 0.02, and the Hardy–Weinberg equilibrium test gave (P>1e^-5^). Remaining missing genotypes were phased and imputed with Beagle v5.4^53^. Additive allele dosages, coded as 0, 1 or 2, were exported with PLINK v1.90b7.2^54^. The final genomic dataset comprised 3,575 individuals and 284,452 SNPs.

### Trait-specific datasets and covariates

For each trait, analyses were restricted to individuals with both phenotype and genotype data. Values below (Q_1_-1.5xIQR) or above (Q_3_+1.5xIQR) were excluded separately within each trait. GWAS sample sizes were 1,274–1,316 for egg-laying traits, 2,605 for EHR, 303–320 for larval-mass traits, 1,249 for PW, 1,580 for AW, 2,432 for DTEP and 308 for DTPA. Generation was included as a fixed effect for reproductive and larval-mass traits. AW and DTEP were adjusted for sex and generation, PW was adjusted for sex and DTEP class, and DTPA was analysed without additional fixed effects. Candidate fixed effects were retained when (*P*<0.05) in preliminary one-way ANOVA for generation.

### Genome-wide association analyses and QTL definition

Each trait was analysed independently with five complementary frequentist approaches. A single-marker linear mixed model was fitted using statgenGWAS v1.0.9^55^:

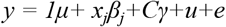

where *x*_*j*_ contained additive genotype dosages for SNP_j_, *β*_*j*_ was its allele-substitution effect, *C* contained the trait-specific fixed effects, and 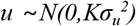. The genomic relationship matrix *K* was calculated from genome-wide markers using the VanRaden method. Variance components were estimated before generalized-least-squares marker testing. Three multi-locus models, MLMM^56^, FarmCPU^57^ and BLINK^58^, were fitted with GAPIT v3.5.0^59^ using the same trait-specific fixed effects. MLMM iteratively fitted associated markers as fixed cofactors in a mixed model. FarmCPU alternated fixed-effect marker tests with random-effect selection of pseudo-quantitative trait nucleotides. BLINK iteratively selected cofactors using linkage disequilibrium and the Bayesian information criterion. Single-step GWAS was performed with the BLUPF90 suite^60^. Genotype and pedigree records were prepared with RENUMF90. Trait-specific additive genetic, common-environment and residual variance components were estimated using average-information REML with a convergence criterion of 10^−12^ and a maximum of 100 iterations. Converged estimates were transferred to the single-step GBLUP and POSTGSF90 parameter files. SNP effects were back-solved from genomic breeding values, and marker P-values were calculated with the snp_p_value option. Only results generated with converged variance-component estimates were retained.

Benjamini–Hochberg correction was applied separately to every trait-by-method scan. Associations with FDR-adjusted (*P*<0.05) were considered significant. SNPs significant in at least two of the five methods were classified as multi-model associations. Genomic inflation was assessed using *λ*_*GC*_, calculated from the median one-degree-of-freedom *χ*^*2*^ statistics, together with quantile–quantile plots. All SNPs significant in at least one method were used to construct candidate QTLs. Genome-wide linkage disequilibrium was estimated as pairwise *r*^*2*^ in PLINK v1.9 using a reproducible 5% sample of 14,077 markers and intrachromosomal marker pairs separated by no more than 2 Mb. Mean *r*^*2*^ first remained below 0.10 in the 25–50-kb distance class; its upper boundary of 50 kb was therefore used as the QTL-grouping distance. Within each trait and chromosome, significant SNPs separated by consecutive gaps of no more than 50 kb were grouped. Each interval was extended by 50 kb on both sides, and overlapping intervals were merged. QTLs belonging to different traits remained distinct. This procedure generated the 305-QTL catalogue used for subsequent comparisons with population-genomic signals.

### QTL overlap with diversity and introgression signals

The 305 QTLs were compared with lower-tail Tajima’s *D* and π regions in the French population and upper-tail *F*_dM_ regions from comparisons in which France was the inferred recipient, using matched 10%, 5% and 1% thresholds. A diversity window was considered farm-specific in the French population when it crossed the relevant lower-tail threshold in France, while the coordinate-matched *Serbia* window was callable and remained above its corresponding threshold. Comparison-specific *F*_dM_ thresholds were estimated from positive values in valid 100-kb windows containing at least 100 SNPs and showing Patterson’s *D* < 0 in France. Each comparison used 100,000 random placements preserving chromosome and QTL width. Random placements were centred on SNPs from the full GWAS scan and matched to each QTL’s callable fraction. After excluding QTLs without callable bases, 282 QTLs were eligible for diversity and joint comparisons and 305 for *F*_dM_ alone. Two-sided empirical *P*-values were calculated from the permutation distributions, and Benjamini–Hochberg correction was applied jointly across all displayed primary French-population tests.

### Software and reproducibility

Analyses were conducted in R v4.3.1 and with custom Python scripts. Principal R packages included poolfstat v3.1.0, statgenGWAS v1.0.9, GAPIT v3.5.0, qgg v1.1.6, CMplot v4.5.1, ggplot2 v4.0.1, data.table v1.17.0, dplyr v1.1.4, tidyr v1.3.1 and stringr v1.5.1. Python-based interval, permutation and visualization analyses used NumPy, pandas, SciPy and Matplotlib. OpenAI’s ChatGPT and Codex, using the GPT-5.6 Sol and GPT-6 Astra models, assisted with coding, debugging and language polishing. All outputs were reviewed and validated by the authors, who take full responsibility for the final code and manuscript. Analysis code and supporting data are publicly available in the TenebrioEvolution GitHub repository: https://github.com/madoui/TenebrioEvolution. The VCF file of pool-seq data is available from Zenodo at https://doi.org/10.5281/zenodo.23099934.

## Acknowledgements

We thank Genoscope, Centre National de Séquençage (CEA, Institut de biologie François-Jacob) for sequencing, the GENTYANE high-throughput genotyping and sequencing platform (INRAE, UMR 1095 GDEC, Clermont-Ferrand) for its technical support, and Yannick Moret and UMR 6282 Biogéosciences (CNRS/Université Bourgogne Europe) for their support and contribution to this work.

## References

1. Larson, G. & Fuller, D. Q. The Evolution of Animal Domesticatio Annu. Rev. Ecol. Evol. Syst. 45, 115–136 (2014).

2. Marshall, F. B., Dobney, K., Denham, T. & Capriles, J. Evaluating the roles of directed breeding and gene flow in animal domestication. Proc. Natl. Acad. Sci. 111, 6153–6158 (2014).

3. Frantz, L. A. F., Bradley, D. G., Larson, G. & Orlando, L. Animal domestication in the era of ancient genomics. Nat. Rev. Genet. 21, 449–460 (2020).

4. Larson, G. & Burger, J. A population genetics view of animal domestication. Trends Genet. 29, 197–205 (2013).

5. Xiang, H et al. The evolutionary road from wild moth to domestic silkworm. Nat. Ecol. Evol. 2, 1268–1279 (2018).

6. Rubin, C.-J et al. Whole-genome resequencing reveals loci under selection during chicken domestication. Nature 464, 587–591 (2010).

7. Girdland Flink, L et al. Establishing the validity of domestication genes using DNA from ancient chickens. Proc. Natl. Acad. Sci. 111, 6184–6189 (2014).

8. Daly, K. G et al. Herded and hunted goat genomes from the dawn of domestication in the Zagros Mountains. Proc. Natl. Acad. Sci. 118, e2100901118 (2021).

9. Daly, K. G et al. Ancient goat genomes reveal mosaic domestication in the Fertile Crescent. Science 361, 85–88 (2018).

10. Frankham, R. Genetic adaptation to captivity in species conservation programs. Mol. Ecol. 17, 325–333 (2008).

11. Barghi, N., Hermisson, J. & Schlötterer, C. Polygenic adaptation unifying framework to understand positive selection. Nat. Rev. Genet. 21, 769–781 (2020).

12. Carneiro, M. et al. Rabbit genome analysis reveals a polygenic basis for phenotypic change during domestication. Science 345, 1074–1079 (2014).

13. Frantz, L. A. et al. Evidence of long-term gene flow and selection during domestication from analyses of Eurasian wild and domestic pig genomes. Nat. Genet. 47, 1141–1148 (2015).

14. Christie, M. R., Marine, M. L., Fox, S. E., French, R. A. & Blouin, M. S. A single generation of domestication heritably alters the expression of hundreds of genes. Nat. Commun. 7, 10676 (2016).

15. Silvaraju, S., Loh, R. K., Kittelmann, S. & Puniamoorthy, N. Genetic differentiation in domesticated black soldier fly (Hermetia illucens) populations. BMC Biol. 24, 7 (2025).

16. Panagiotakopulu, E. New Records for Ancient Pests: Archaeoentomology in Egypt. J. Archaeol. Sci. 28, 1235–1246 (2001).

17. Pals, J. P. & Hakbijl, T. Weed and insect infestation of a grain cargo in a ship at the Roman fort of Laurium in Woerden (Province of Zuid-Holland). Rev. Palaeobot. Palynol. 73, 287–300 (1992).

18. Bain, A., Bouchard-Perron, J.-A., Auger, R. & Simoneau, D. Bugs, seeds and weeds at the Intendant’s Palace: a study of an evolving landscape. Post-Mediev. Archaeol. 43, 183–197 (2009).

19. Marba. Made in Apeldoorn https://madeinapeldoorn.com/english/marba-3/.

20. Cotton, R. T. Notes on the Biology of the Meal Worms, Tenebrio Molitor Linne and T. Obscurus Fab. Ann. Entomol. Soc. Am. 20, 81–86 (1927).

21. Morales-Ramos, J. A., Tomberlin, J. K., Miranda, C. & Rojas, M. G. Rearing methods of four insect species intended as feed, food, and food ingredients: a review. J. Econ. Entomol. 117, 1210–1224 (2024).

22. Morales-Ramos, J. A. & Rojas, M. G. Effect of Larval Density on Food Utilization Efficiency of Tenebrio molitor (Coleoptera: Tenebrionidae). J. Econ. Entomol. 108, 2259–2267 (2015).

23. Lenormand, T. Gene flow and the limits to natural selection. Trends Ecol. Evol. 17, 183–189 (2002).

24. Kaya, C. et al. Global population genetic structure and demographic trajectories of the black soldier fly, Hermetia illucens. BMC Biol. 19, 94 (2021).

25. House, J. Insects as food in the Netherlands: Production networks and the geographies of edibility. Geoforum 94, 82–93 (2018).

26. Orozco-terWengel, P. et al. Adaptation of Drosophila to a novel laboratory environment reveals temporally heterogeneous trajectories of selected alleles. Mol. Ecol. 21, 4931–4941 (2012).

27. Christie, M. R., Marine, M. L., French, R. A. & Blouin, M. S. Genetic adaptation to captivity can occur in a single generation. Proc. Natl. Acad. Sci. 109, 238–242 (2012).

28. Deruytter, D., Coudron, C. L. & Claeys, J. The Effects of Density on the Growth and Temperature Production of Tenebrio molitor Larvae. Sustainability 14, 6234 (2022).

29. van Broekhoven, S., Oonincx, D. G. A. B., van Huis, A. & van Loon, J. J. A. Growth performance and feed conversion efficiency of three edible mealworm species (Coleoptera: Tenebrionidae) on diets composed of organic by-products. J. Insect Physiol. 73, 1–10 (2015).

30. Barnes, A. I. & Siva-Jothy, M. T. Density–dependent prophylaxis in the mealworm beetle Tenebrio molitor L. (Coleoptera: Tenebrionidae): cuticular melanization is an indicator of investment in immunity. Proc. R. Soc. B Biol. Sci. 267, 177–182 (2000).

31. Slocinska, M., Mirek, J., Adamski, Z. & Lubawy, J. The Activity of Solanum tuberosum Leaf Extract and Chaconine in the Gut of Tenebrio molitor Larvae. Toxins 18, 157 (2026).

32. Chipoulet, J.-M. & Chararas, C. Survey and electrophoretical separation of the glycosidases of Rhagium inquisitor (Coleoptera: Cerambycidae) larvae. Comp. Biochem. Physiol. Part B Comp. Biochem. 80, 241–246 (1985).

33. Barghi, N. et al. Genetic redundancy fuels polygenic adaptation in Drosophila. PLOS Biol. 17, e3000128 (2019).

34. Coudron, C. L. et al. BugBook: Basic information and good practices on how to maintain stock populations for Tenebrio molitor and Hermetia illucens for research. J. Insects Food Feed 11, 241–267 (2025).

35. Morales-Ramos, J. A., Kelstrup, H. C., Rojas, M. G. & Emery, V. Body Mass Increase Induced by Eight Years of Artificial Selection in the Yellow Mealworm (Coleoptera: Tenebrionidae) and Life History Trade-offs. J. Insect Sci. 19, 4 (2019).

36. Sellem, E. et al. Multitrait genetic parameter estimates in a Tenebrio molitor reference population: high potential for breeding gains. animal 18, 101197 (2024).

37. Wu, D. et al. Pervasive introgression facilitated domestication and adaptation in the Bos species complex. Nat. Ecol. Evol. 2, 1139–1145 (2018).

38. Hermisson, J. & Pennings, P. S. Soft Sweeps: Molecular Population Genetics of Adaptation From Standing Genetic Variation. Genetics 169, 2335–2352 (2005).

39. Zhang, Y. et al. The octopamine receptor, OA2B2, modulates stress resistance and reproduction in Nilaparvata lugens Stål (Hemiptera: Delphacidae). Insect Mol. Biol. 31, 33–48 (2022).

40. Orlando, L. et al. Ancient DNA analysis. Nat. Rev. Methods Primer 1, 14 (2021).

41. Orlando, L., Gilbert, M. T. P. & Willerslev, E. Reconstructing ancient genomes and epigenomes. Nat. Rev. Genet. 16, 395–408 (2015).

42. Li, H. Aligning sequence reads, clone sequences and assembly contigs with BWA-MEM. (2013).

43. Li, H. et al. The Sequence Alignment/Map format and SAMtools. Bioinforma. Oxf. Engl. 25, 2078–2079 (2009).

44. Margarido, G. R. A., Souza, A. P. & Garcia, A. a. F. OneM software for genetic mapping in outcrossing species. Hereditas 144, 78–79 (2007).

45. Tang, H. et al. ALLMAPS: robust scaffold ordering based on multiple maps. Genome Biol. 16, 3 (2015).

46. Chen, S. fastp 1.0: An ultra-fast all-round tool for FASTQ data quality control and preprocessing. iMeta 4, e70078 (2025).

47. Gautier, M., Vitalis, R., Flori, L. & Estoup, A. f-Statistics estimation and admixture graph construction with Pool-Seq or allele count data using the R package poolfstat. Mol. Ecol. Resour. 22, 1394–1416 (2022).

48. Lê, S., Josse, J. & Husson, F. FactoMineR: An R Package Multivariate Analysis. J Stat. Softw. 25, 1–18 (2008).

49. Kofler, R., Pandey, R. V. & Schlötterer, C. PoPoolatio Identifying differentiation between populations using sequencing of pooled DNA samples (Pool-Seq). Bioinformatics 27, 3435–3436 (2011).

50. Ashburner, M. et al. Gene ontology: tool for the unification of biology. The Gene Ontology Consortium. Nat. Genet. 25, 25–29 (2000).

51. Cingolani, P. et al. A program for annotating and predicting the effects of single nucleotide polymorphisms, SnpEff. Fly (Austin) 6, 80–92 (2012).

52. Benjamini, Y. & Hochberg, Y. Controlling the False Discovery Rate: A Practical and Powerful Approach to Multiple Testing. J. R. Stat. Soc. Ser. B Methodol. 57, 289–300 (1995).

53. Browning, B. L., Zhou, Y. & Browning, S. R. A One-Penny Imputed Genome from Next-Generation Reference Panels. Am. J. Hum. Genet. 103, 338–348 (2018).

54. Chang, C. et al. Second-generation PLINK: Rising to the challenge of larger and richer datasets. GigaScience 4, 1–16 (2015).

55. Rossum, B.-J. van & Kruijer,W. statgenGWAS: Genome Wide Association Studies. (2022).

56. Segura, V. et al. An efficient multi-locus mixed model approach for genome-wide association studies in structured populations. Nat. Genet. 44, 825–830 (2012).

57. Kusmec, A. & Schnable, P. S. FarmCPUpp: Efficient large-scale genomewide association studies. Plant Direct 2, e00053 (2018).

58. Huang, M., Liu, X., Zhou, Y., Summers, R. M. & Zhang, Z. BLINK: a package for the next level of genome-wide association studies with both individuals and markers in the millions. GigaScience 8, giy154 (2019).

59. Lipka, A. et al. GAPIT: genome association and prediction integrated tool. Bioinformatics 28, 2397–2399 (2012).

60. Aguilar, I et al. BLUPF90 Suite of Programs for Animal Breeding with Focus on Genomics. (2018).

