## Extended Data for "Mosaic selection and gene flow at the onset of grain beetle domestication"


Lucas Rocha Ferreira[1], Romuald Laso-Jadart[2], Leonardo Panunzi[1], Sana Haddad[1], Evangelia Eleftheriou[3], Benoit Vacherie[3], Eliaou Sellem[4,5], Karine Labadie[3], Thomas Lefebvre[5], Christophe Ambroise[6], Mohammed-Amin Madoui[1]

[1]Service d'Etude des Prions et des Infections Atypiques (SEPIA), Institut François Jacob, Commissariat à l'Energie Atomique et aux Énergies Alternatives (CEA), Université Paris Saclay, Fontenay-aux-Roses, France

[2]Univ Paris-Est Créteil, Sorbonne Université, Université Paris- Cité, CNRS, IRD, INRAE, Institute of Ecology and Environmental Science, IEES, Créteil, France

[3]Genoscope, Institut de biologie François Jacob, CEA, Université Paris-Saclay, 91057 Evry, France

[4]UVSQ, INRAE, BREED, Université Paris-Saclay, Jouy-en-Josas, France

[5]Ynsect, 91000, Evry, France

[6]Laboratoire de Mathématiques et Modélisation d'Évry, Université Paris-Saclay, CNRS, Univ Evry


##### Extended Data Figure 1: Study overview

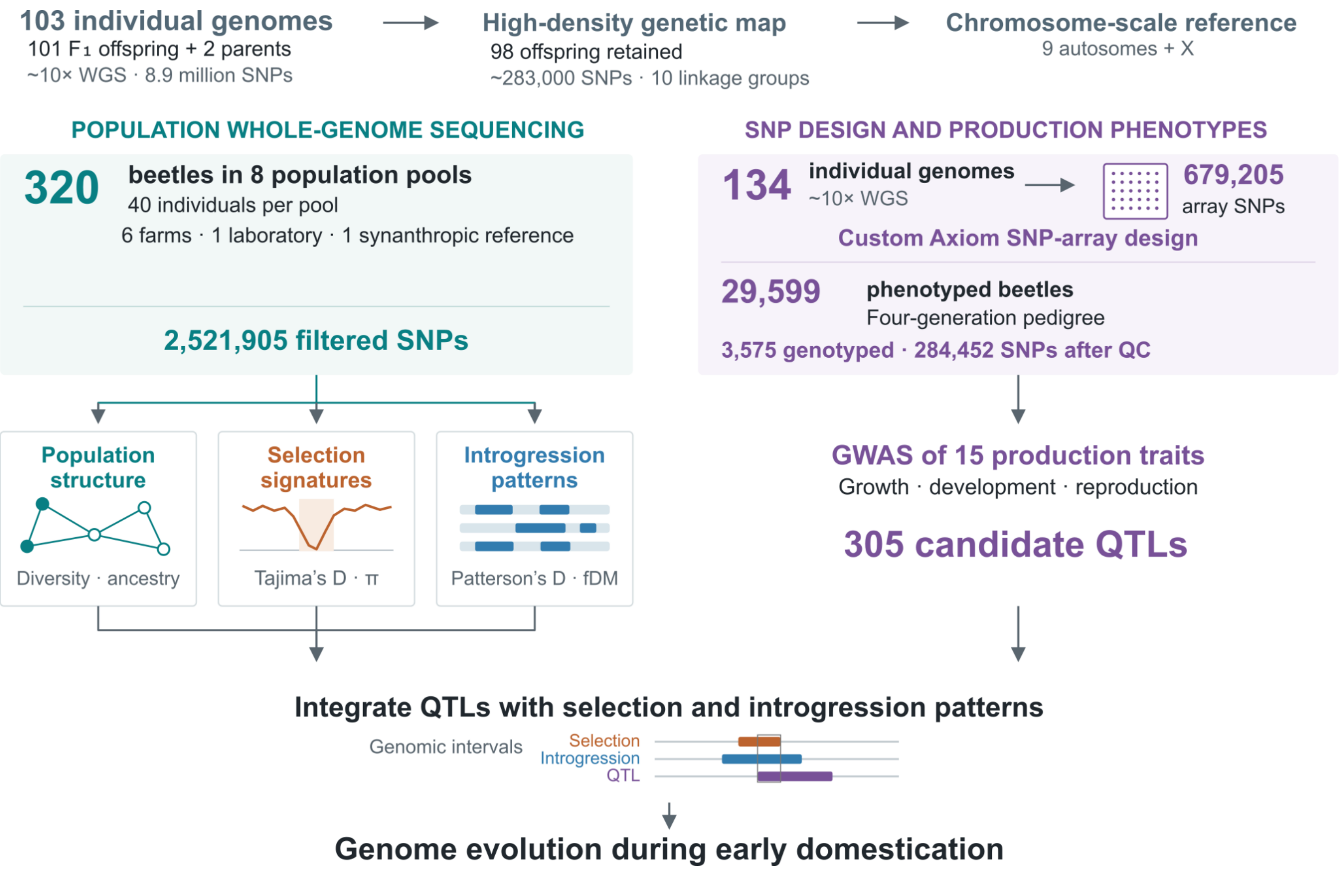


Overview of the genomic resources and analyses used to investigate early domestication in *Tenebrio molitor*. Whole-genome sequencing of two parents and 101 $F_1$ offspring supported construction of a genetic map and a chromosome-scale reference comprising nine autosomes and the X chromosome; 98 offspring were retained for mapping. Population genomic analyses used pooled sequencing of 320 beetles from eight populations, with 40 individuals per pool, representing six farms, one laboratory population and one synanthropic reference. Separately, sequencing of 134 individuals supported SNP-array design. A four-generation pedigree comprising 29,599 phenotyped beetles included 3,575 genotyped individuals, with 284,452 SNPs retained after quality control. Genome-wide association analyses of 15 production traits identified 305 QTLs. Genomic overlap analyses compared these QTLs with candidate selection and introgression signals.

#### Extended Data Figure 2: Population contributions to regional $F_{ST}$ outliers.

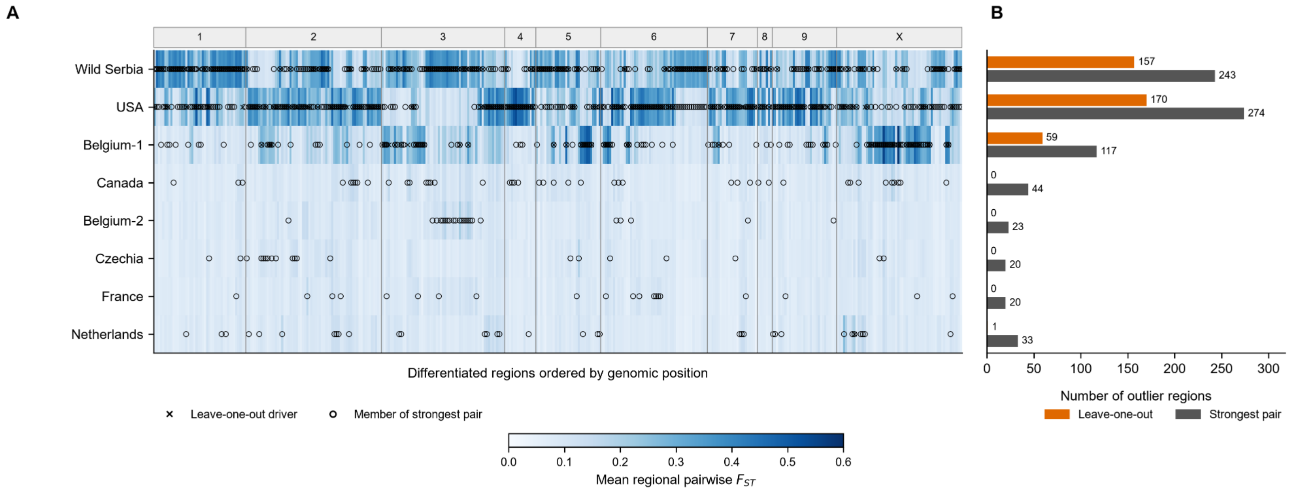


**A**, Mean pairwise FST for each population across 387 regions obtained by merging overlapping 50-SNP windows exceeding the empirical 99th percentile of multi-population $F_{ST}$ ($F_{ST}$ > 0.156857; upper 1% tail). Rows represent populations and columns represent regions ordered by genomic position; the upper strip identifies chromosomes. Crosses mark the population whose exclusion caused the largest decrease in regional multi-population $F_{ST}$. Open circles identify the two members of the strongest regional pairwise contrast. **B**, Numbers of regions assigned to each population as the leave-one-out driver (orange) or as a member of the strongest pair (grey). Each region contributes one driver assignment and two strongest-pair memberships.

#### Extended Data Table 1: Admixture graph summary metrics

| Metric | Jackknife block size (SNPs) | | | | |
|---|---|---|---|---|---|
| | 1,000 | 5,000 | 10,000 | 20,000 | 40,000 |
| Jackknife blocks | 2,507 | 495 | 243 | **116** | 53 |
| Admixture edges | 3 | 3 | 3 | **3** | 2 |
| BIC | 590.08 | 550.68 | 544.20 | **541.35** | 552.52 |
| Optimizer convergence code | 52 | 0 | 0 | **0** | 0 |
| Maximum \|residual Z\| | 3.46 | 2.74 | 2.71 | **2.03** | 2.83 |
| Residuals with \|Z\| > 2 | 34 | 7 | 4 | **1** | 36 |
| Residuals with \|Z\| > 3 | 7 | 0 | 0 | **0** | 0 |
| Final candidate graphs | 28 | 15 | 23 | **14** | 38 |
| Graphs with ΔBIC < 2 | 8 | 2 | 4 | **3** | 2 |
| Graphs with ΔBIC < 6 | 28 | 15 | 23 | **14** | 38 |
| ΔBIC to next-best graph | $2.18 \times 10^{-11}$ | 1.59 | 0.52 | **0.34** | 1.14 |

Values summarize the admixture graphs shown in Extended Data Figure 3. Bold values identify the 20,000-SNP analysis retained for the principal analysis in the manuscript. Optimizer convergence codes: 0 indicates successful completion; 52 indicates an L-BFGS-B optimizer error. The 1,000-SNP result therefore did not converge successfully and is retained here as a flagged diagnostic result. ΔBIC is the difference from the lowest BIC among graphs fitted within the same block-size analysis. The next-best graph is the graph with the second-lowest BIC. Counts of graphs within each ΔBIC threshold refer to the final retained set. Absolute BIC values are not used to rank block sizes, because changing the block size changes the estimated covariance structure used for fitting. Residual Z denotes a standardized graph residual. The maximum absolute residual and the numbers exceeding |Z| = 2 or 3 summarize graph misfit. The 20,000-SNP analysis has the smallest maximum absolute residual among the analyses shown, but retains three graphs within ΔBIC < 2; its next-best graph differs by only 0.34 BIC units.

**Extended Data Figure 3: Admixture graphs with different number of SNPs per block size**

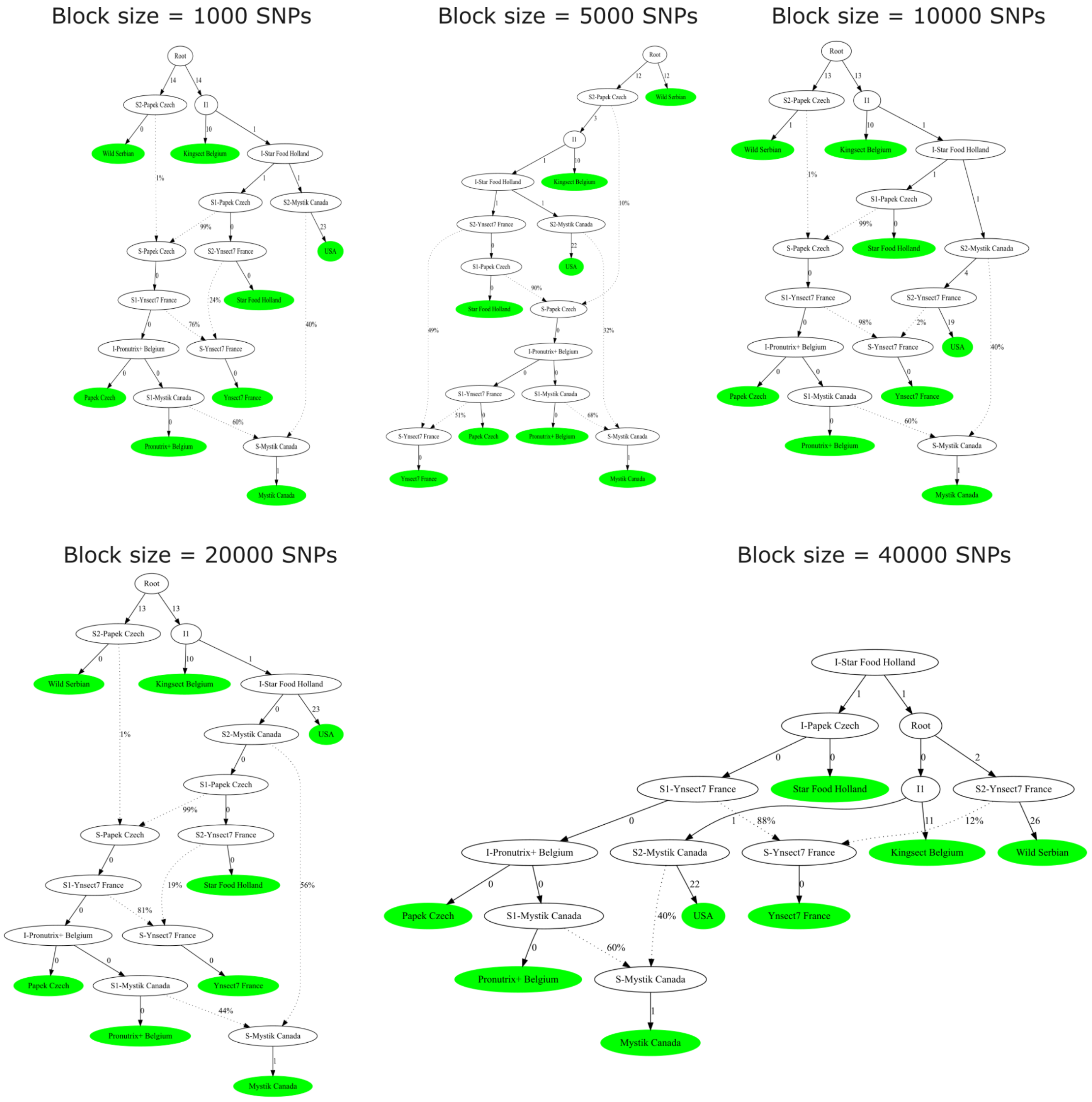


Admixture graphs inferred from population allele-frequency statistics using jackknife blocks of 1,000, 5,000, 10,000, 20,000 and 40,000 SNPs. Green nodes represent sampled populations, whereas white nodes represent inferred ancestral lineages. Solid branches represent genetic drift, with adjacent numbers indicating fitted drift parameters. Dotted arrows represent inferred admixture contributions, and percentages indicate the estimated contribution of each parental lineage to the corresponding admixed lineage. Each panel shows the graph obtained using the indicated block size. The graph inferred with 20,000-SNP blocks was retained for the principal analysis. Corresponding graph-fit statistics are reported in Extended Data Table 1.

#### Extended Data Figure 4: Introgression analysis.

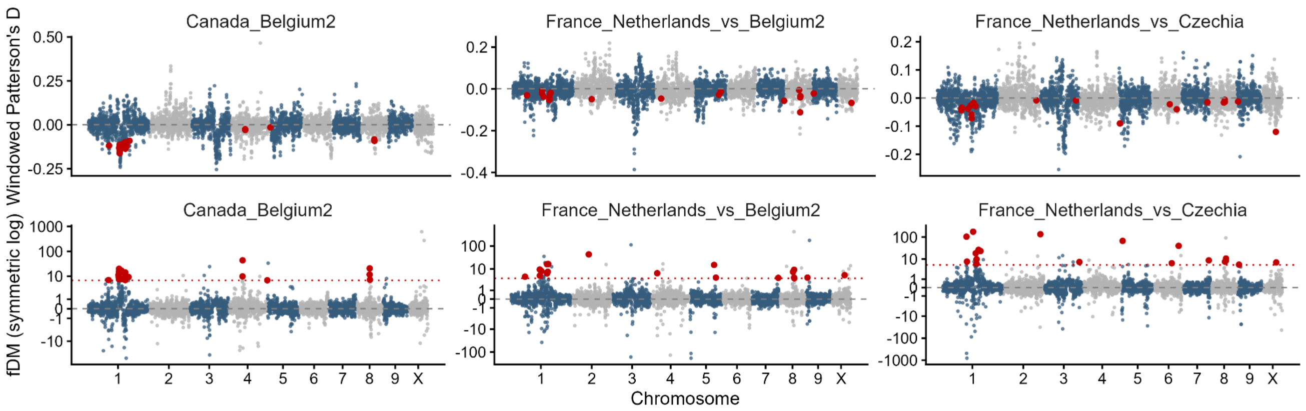


Genome-wide Patterson's $D$ (top) and $F_{dM}$ (bottom) calculated in 100-kb windows. Chromosomes alternate blue and grey; stringent upper-1% $F_{dM}$ candidates with the expected Patterson's $D$ direction are red.

#### Extended Data Figure 5: Distributions $p_N/p_S$ log ratio of low Tajima'*D* loci versus background in farmed populations

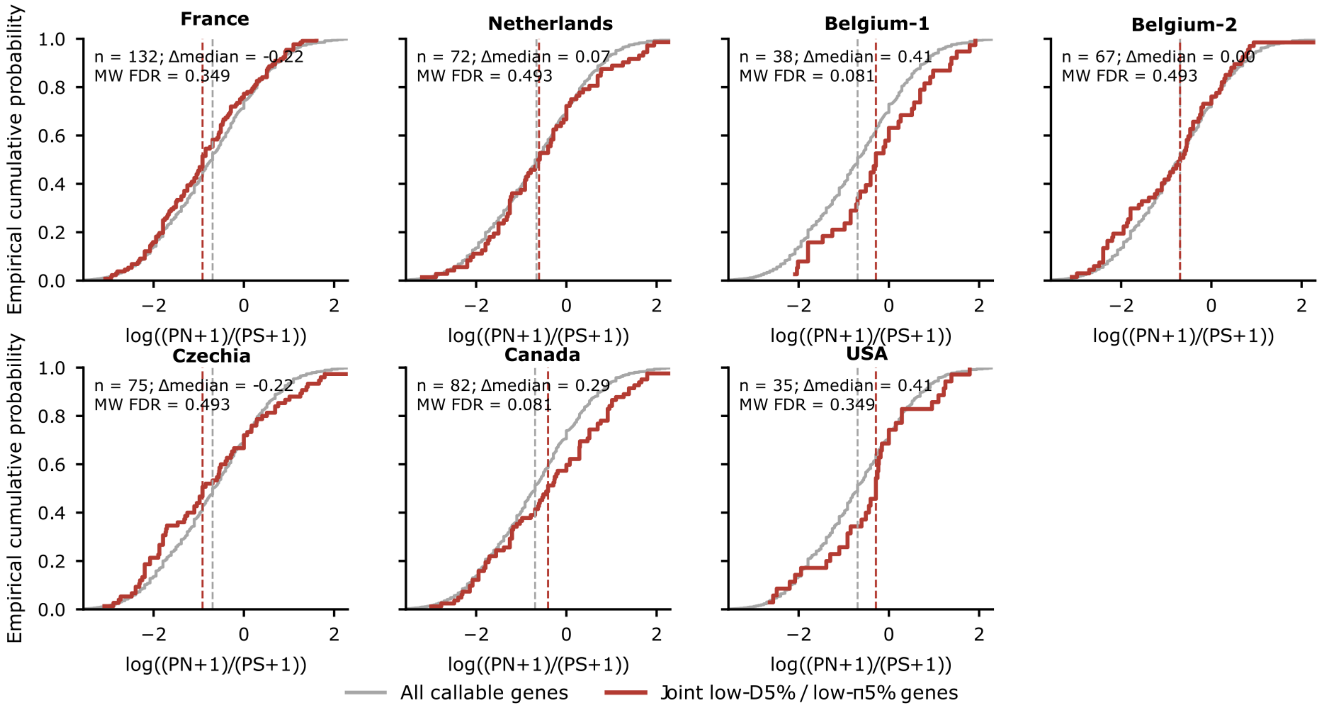


Empirical cumulative distributions of $\ln[(P_n + 1)/(P_s + 1)]$, where $P_n$ and $P_s$ are the numbers of segregating nonsynonymous and synonymous coding variants, respectively. Analyses included genes with at least five coding variants and callable diversity estimates. Grey curves show all eligible genes; red curves show focal genes meeting the population-specific lower-5% criteria for both Tajima's D and nucleotide diversity (π). Panels represent the six farm populations and the USA laboratory population. Dashed vertical lines indicate the medians of the corresponding plotted distributions. Panel annotations report the number of focal genes (n), the focal-minus-nonfocal median difference (Δmedian), and the two-sided Mann–Whitney test P value adjusted using the Benjamini–Hochberg procedure across the seven populations (MW FDR). Statistical comparisons use focal versus nonfocal genes, whereas the grey curves include both groups.

### Extended Data Note 1: GEBV and QTL analysis

#### Genomic prediction of development time from egg to pupa

We next asked whether DTEP-associated regions identified by GWAS retained explanatory value in an independent validation population. Three marker sets were compared: a whole-genome panel of 3,554,876 SNPs, a robust-QTL panel of 53,854 SNPs located within robust DTEP QTL intervals, and a reduced panel of 77 significant SNPs. In GBLUP models fitted in the validation cohort, the whole-genome panel showed the highest explanatory capacity for DTEP ($h^2_{SNP} = 0.58$; $R^2 = 0.87$). The robust-QTL panel, despite representing only 1.5% of the genome-wide marker set, retained substantial signal ($h^2_{SNP} = 0.21$; $R^2 = 0.38$), whereas the 77-SNP panel captured a smaller but still detectable fraction of phenotypic variance ($h^2_{SNP} = 0.12$; $R^2 = 0.2$). These results indicate that robust DTEP QTL intervals contain reproducible genomic information beyond the GWAS discovery population and provide a first proof of concept for QTL-guided marker-panel development in *T. molitor*.

#### Analysis of DTEP QTL candidates

The 2,432 individuals had a development time from egg to pupa (DTEP) median of 77 days (Extended Data Fig. 6a), and were classified into four categories: fast, fast-moderate, slow-moderate, and slow. Bayesian GWAS used BSLMM in GEMMA v0.96 and BayesB, BayesC and BayesR in GCTB v2.04.3. BSLMM combines sparse SNP effects with a polygenic background, whereas the other models use alternative mixture priors for SNP effects. Posterior inclusion probabilities (PIP) were estimated from post-burn-in MCMC samples, with PIP $\geq$ 0.4 used to retain candidate SNPs. Across the four Bayesian models, the mean heritability estimate was 0.37 ± 0.05. Using five frequentist and four Bayesian GWAS methods, we identified 75 candidate genomic regions after merging overlapping QTLs and grouping associated signals according to linkage disequilibrium, whereas the main analysis used 88 DTEP QTLs defined using the 50-kb grouping criterion (Extended Data Fig. 6b-c). Nine genomic regions supported by at least two methods were classified as robust QTLs and three genomic regions supported by at least one frequentist method (FDR < 0.05) and one Bayesian method (PIP > 0.4) were classified as very robust DTEP-6, DTEP-8 (Extended Data Fig. 6e-h), and DTEP-9 . Ontology term enrichment analysis of the 281 genes within DTEP QTLs showed four molecular function terms significantly enriched (FDR = 0.05), and were all related to cuticle structure, making cuticle-related molecular functions the only robust enrichment signal for DTEP. Three QTLs (DTEP-1, 6 and 8) had the strongest combination of statistical support and interpretable candidate genes (Extended Data Fig. 6e-h).

DTEP-1 was a broad chromosome 2 locus spanning 1.41 Mb, supported by three frequentist models and containing 164 significant SNPs, including 68 robust SNPs. The lead marker AX-677863924 showed strong support (BLINK $-\log_{10}P$ = 13.37, $P \approx 4.3 \times 10^{-14}$; mean frequentist $-\log_{10}P$ = 9.44) and marked an additive shift in DTEP classes, with the alternative allele associated with faster development. The strongest coding candidate was a FIRMM-like gene intersected by 28 significant SNPs, including 22 robust SNPs and 13 coding variants, suggesting a link with genome maintenance, DNA repair or replication-associated developmental processes. DTEP-6 was a compact 108.73 kb locus with 25 significant SNPs, 32 LD markers and 0.45–0.89% variance explained. The lead SNP (AX-674418014) was supported by MLM, BLINK and BayesB, with a maximum $-\log_{10}P$ = 10.95 , mean $-\log_{10}P$ = 4.48 and maximum PIP = 0.549. Genotype classes showed an additive pattern, with the alternative allele associated with slower development. The signal centred on a LOTUS–Tudor gene, and nearby JH-esterase-like carboxylesterases, suggesting a candidate module linking maternal RNA regulation, early development and endocrine timing of larval–pupal transition. DTEP-8 was a very robust 102.94 kb locus supported by BLINK and four Bayesian models, with four significant SNPs and 0.69–1.20% variance explained. The lead missense SNP (AX-675302864; Thr23Met) mapped to an ankyrin-repeat protein within a small ankyrin-repeat gene cluster, and showed strong statistical support (BLINK $-\log_{10}P$ = 13.62; mean PIP = 0.722). Unlike the other two loci, DTEP-8 showed a non-additive pattern consistent with underdominance: both homozygous genotypes were enriched for faster development, whereas heterozygous individuals were shifted toward slower developmental classes.

#### Analysis of EHR QTL candidates

Egg hatching rate (EHR) distribution across 2,605 individuals had a median of 79.03% (Extended Data Fig. 7a) and a mean heritability estimate of 0.143 ± 0.019 across the four Bayesian models. Applying the same merging and QTL prioritization used for DTEP, we identified nine robust QTLs, and four very robust QTLs (Extended Data Fig. 7b). GO terms enrichment analysis of genes within QTLs showed 28 significantly enriched biological processes (FDR<0.05), 23 mapped to a compact gustatory receptor locus, and the two molecular function terms were taste receptor activity and sweet taste receptor activity. Among the nine robust QTLs, three combined strong statistical support with interpretable candidate genes or variant annotations (EHR-3, 4 and 8). EHR-3 on chromosome 5 encompassed 69 significant SNPs across approximately 0.99 Mb. Its lead SNP, AX-676384541, was intronic within a Pgant35A-like polypeptide N-acetylgalactosaminyltransferase gene and reached a maximum $-\log_{10} P$ of 12.19. EHR-4, also on chromosome 5, spanned 1.54 Mb and contained 29 significant SNPs. Its lead SNP, AX-676901494, was a missense variant predicting a Thr134Ala substitution in a hemocyte transglutaminase-like protein, with a mean $-\log_{10} P$ of 6.99 across the supporting MLM and MLMM models. EHR-8 on chromosome 8 showed the broadest support, with associations detected across all nine GWAS models. This locus contained two intronic SNPs within a gene encoding a Tubby C-terminal domain-containing protein. AX-681166815 was supported by all five frequentist models and reached a maximum $-\log_{10} P$ of 11.80, whereas the neighbouring SNP AX-681144307 was supported by four models and reached 15.66.

#### Extended Data Figure 6: QTLs results for GWAS analysis on DTEP

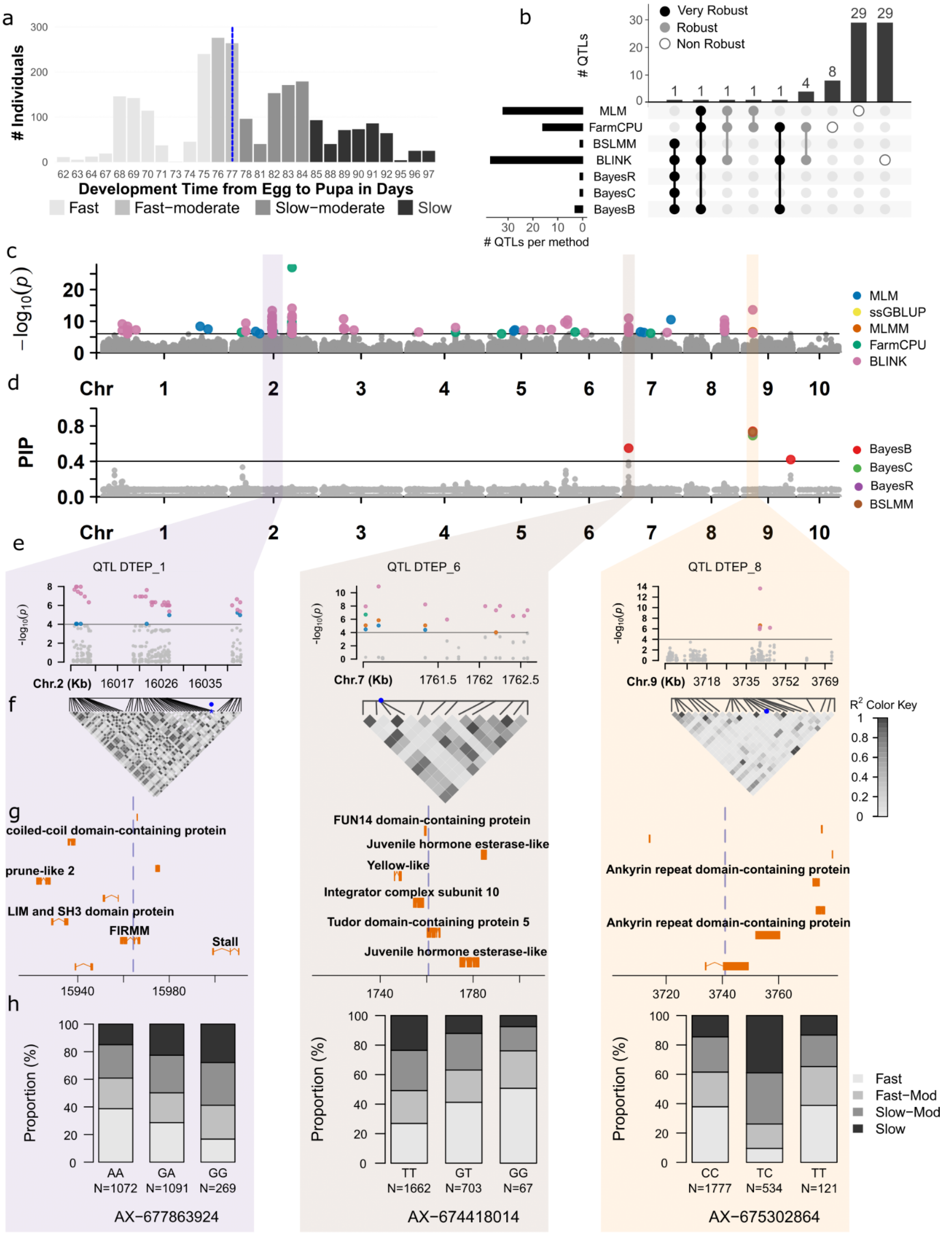


**(a)** Distribution of DTEP across individuals, expressed in days. Shading distinguishes four development classes: fast, fast–moderate, slow–moderate, and slow. The blue dashed line marks 77 days. **(b)** UpSet plot summarizing QTL detection across seven analytical methods. Horizontal bars indicate the number of QTLs detected by each method, and vertical bars indicate the number detected by the corresponding combinations of methods. Black, grey, and open circles denote very robust, robust, and non-robust detections, respectively. **(c, d)** Genome-wide distributions of association statistics ($-\log_{10} P$) and posterior inclusion probabilities (PIP) across the ten chromosomes. Colored points identify results from the methods indicated in the legends. Horizontal lines mark the association threshold in **(c)** and PIP = 0.4 in **(d)**. Shaded regions highlight QTLs shown in detail below. **(e–h)** Regional characterization of QTL DTEP_1, QTL DTEP_6, and QTL DTEP_8 on chromosomes 2, 7, and 9, respectively, comprising **(e)** regional association plots, **(f)** pairwise linkage disequilibrium heatmaps, **(g)** annotated gene models, and **(h)** genotype-specific proportions of development classes. In **(f)**, shading represents linkage disequilibrium ($r^2$), with darker shades indicating higher values. In **(g)**, blue dashed lines indicate focal SNP positions, and genomic coordinates are expressed in kilobases. In **(h)**, stacked bars show the percentage of individuals belonging to each development class within each genotype at SNPs AX-677863924, AX-674418014, and AX-675302864, respectively. Genotype sample sizes (*N*) are indicated below the bars.

**Extended Data Figure 7: QTLs results for GWAS analysis on EHR**

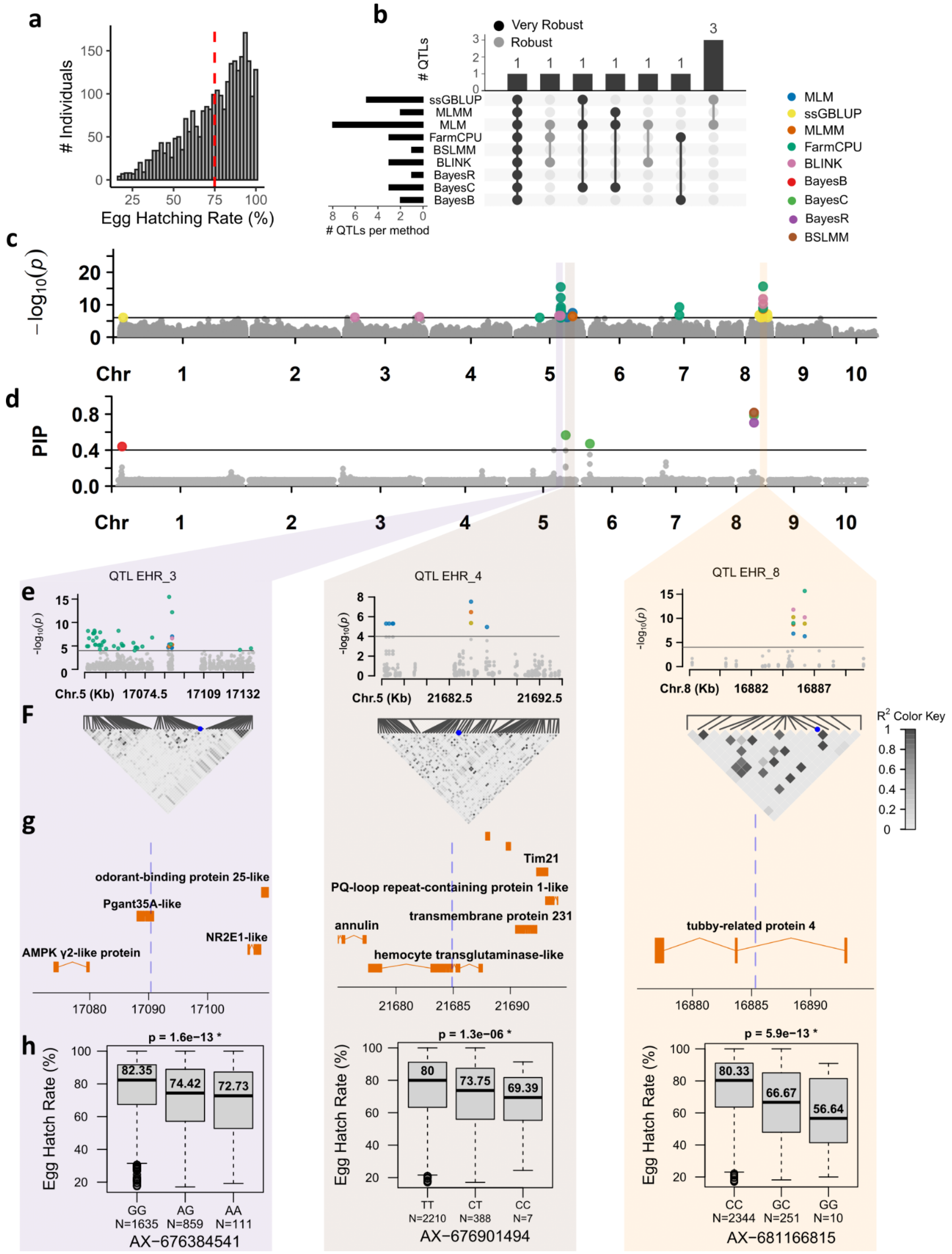


**(a)** Distribution of EHR across individuals; the red dashed line is mean EHR. **(b)** UpSet plot summarizing QTL detection across nine analytical methods. Horizontal bars indicate the number of QTLs detected by each method, and vertical bars indicate the number detected by the corresponding combinations of methods. Black and grey circles denote detections classified as very robust and robust, respectively. **(c, d)** Genome-wide distributions of association statistics ($-\log_{10} P$) and posterior inclusion probabilities (PIP) across the ten chromosomes. Colored points identify results from the methods indicated in the legend. Horizontal lines mark the association threshold in **(c)** and PIP = 0.4 in **(d)**. Shaded regions highlight QTLs shown in detail below. **(e–h)** Regional characterization of QTL EHR_3 and QTL EHR_4 on chromosome 5 and QTL EHR_8 on chromosome 8, comprising **(e)** regional association plots, **(f)** pairwise linkage disequilibrium heatmaps, **(g)** annotated gene models, and **(h)** genotype-specific EHR distributions. In **(f)**, shading represents linkage disequilibrium ($r^2$), with darker shades indicating higher values. In **(g)**, blue dashed lines indicate focal SNP positions, and genomic coordinates are expressed in kilobases. In **(h)**, boxplots show EHR by genotype at SNPs AX-676384541, AX-676901494, and AX-681166815, respectively; genotype sample sizes (*N*) and *P* values are indicated.

**Extended Data Table 2: European Nucleotide Archive (ENA) accession numbers for the sequencing datasets used in this study.**

| Dataset | Biological material | BioProject accession | ENA study accession | ENA run accessions |
|---|---|---|---|---|
| F1 linkage mapping | 103 individuals: 101 F1 offspring and both parents | PRJEB128050 | ERP207507 | ERR18026472–ERR18026574 (103 runs) |
| Population genomics | 320 individuals from eight populations; one DNA pool of 40 individuals per population | PRJEB128051 | ERP207508 | Pending |
| Axiom SNP-array development | 134 individually sequenced members of the pedigreed breeding population | PRJEB128052 | ERP207509 | ERR18026198–ERR18026465 (268 runs) |
